# Benchmarking scRNA-seq copy number variation callers

**DOI:** 10.1101/2024.12.18.629083

**Authors:** Katharina T. Schmid, Aikaterini Symeonidi, Dmytro Hlushchenko, Maria L. Richter, Maria Colomé-Tatché

## Abstract

Copy number variations (CNVs), the gain or loss of genomic regions, are associated with different diseases and cancer types, where they are related to tumor progression and treatment outcome. Single cell technologies offer new possibilities to measure CNVs in individual cells, allowing to assess population heterogeneity and to delineate subclonal structures. Single cell whole-genome sequencing is considered the gold-standard for the quantification of CNVs in single cells. However, the majority of existing single cell datasets interrogate gene expression, using scRNA-seq. Consequently, several computational approaches have been developed to identify CNVs from that data modality. Nevertheless, an independent benchmarking of these methods is lacking. We used 15 scRNA-seq datasets and evaluated six popular computational methods in their ability to recover the ground truth CNVs using a large set of performance metrics. Additionally, we explored whether they could correctly identify euploid cells, especially also in fully diploid samples, and subclonal structures in heterogeneous tumor samples. We discovered several dataset-specific factors that influence the performance of the methods, such as the dataset size and the number and type of CNVs in the analyzed sample. We found that the choice of the reference dataset can have a large impact on the performance. Methods which included additional allelic information from the scRNA-seq reads performed more robustly across scenarios, but at the cost of higher runtime. Furthermore, the methods differed substantially in their additional functionalities and resource requirements. We offer a benchmarking pipeline to help identify the optimal CNV calling method for newly generated scRNA-seq datasets, and to benchmark and improve new methods performance.

## Introduction

Copy number variations (CNVs) describe the gain or loss of genomic regions, from small sequences up to complete chromosomes. These genomic alterations lead to aneuploidy and are associated with different diseases and cancer types^1^. Specific CNVs are hallmarks for the classification of tumors, and are related to tumor progression and treatment outcomes^2–4^. However, the direct functional consequences of CNVs are yet not fully understood^4^. Tumors are very heterogeneous, with different tumor cells having distinct molecular phenotypes. This also applies to CNVs, which can differ substantially between cellular subclones among samples and inside of the same sample, emphasizing the importance of cell-specific analyses^5–7^. Single cell whole-genome sequencing is considered the gold-standard technique to obtain per-cell CNV profiles^8^, as changes in DNA copy number should lead to observable changes in the read sequencing depth. However, the technology is not frequently used in the laboratory compared to other single cell technologies.

Instead, computational methods have been developed to infer the CNV profiles from single cell RNA-seq data^5,9–14^ and single cell assay for transposase-accessible chromatin with sequencing (scATAC-seq) data^15,16^. These approaches have the advantage that apart from the copy number gains and losses, also the information about the cellular state can be obtained from the same measurement (gene expression or open chromatin). For scATAC-seq, the read-out is relatively similar to whole-genome sequencing, as also the genome is sequenced and therefore the read coverage provides information about ploidy. However, for scRNA-seq the inference of CNVs is challenging, as the expression level of genes is highly affected by regulatory mechanisms, and therefore it provides only indirect information about the CNV state. Nevertheless, the general assumption of all computational methods that infer CNVs from scRNA-seq data is that genes in gained regions show higher expression, and in lost regions lower expression, compared to genes in diploid regions. This requires for all methods sophisticated data normalization strategies, using generally reference diploid samples, often in combination with denoising approaches, before the different CNV inference strategies can be applied.

Because of the wealth of scRNA-seq data available, correct identification of CNVs from this data modality is crucial to study the role of CNVs in cancer and other aneuploid tissues. scRNA-seq CNV callers are currently used in many applications, e.g. ^17–20^. However, there is no independent validation that shows whether scRNA-seq CNV calling methods can correctly identify CNVs, and which of the CNV callers works the best.

In this work, we evaluated the performance of six popular CNV callers for scRNA-seq data using 15 different datasets. We evaluated the general CNV prediction performance for each method, comparing its results to a ground truth provided by an orthogonal CNV measurement (either (single cell) whole genome sequencing ((sc)WGS) or whole exome sequencing (WES)), using correlation, area under the curve (AUC) values and F1 scores. Additional aspects included in our evaluation are the prediction results on a diploid sample, the correctness of the inferred clonal structure, the impact of the selected reference dataset for the performance, and the runtime and memory requirements. In addition, we also evaluated the automatic identification of cancer cells for the methods that allow it. Our evaluation is publicly available with a reproducible snakemake pipeline (https://github.com/colomemaria/benchmark_scrnaseq_cnv_callers). This allows the direct testing of new datasets to determine optimal CNV calling strategies, and it facilitates comparisons between methods to improve the performance of newly developed computational tools.

## Results

### scRNA-seq CNV calling benchmarking

We included in our benchmarking study six CNV calling methods that were developed specifically for scRNA-seq data (**Table 1**). The methods can be broadly classified into two categories: one class that uses only the expression levels per gene, consisting of InferCNV^5^, copyKat^12^, SCEVAN^13^ and CONICSmat^9^; and a second class that combines the expression values with minor allele frequency information, consisting of CaSpER^11^ and Numbat^14^. CaSpER and Numbat use allele frequencies per SNP called directly from the scRNA-seq reads, and both models implement a Hidden Markov Model (HMM) to call CNVs. Also InferCNV identifies CNVs using an HMM, but based on expression levels only. copyKat and SCEVAN both apply a segmentation approach, while CONICSmat estimates the CNVs based on a Mixture Model (see **Supplementary Methods**). All methods were run as recommended in the respective tutorials or based on default parameters.

**Table 1.** CNV calling methods from scRNA-seq. HMM = Hidden Markov Model, MM = Mixture Mode, (B)AF = (B-)allele frequency.

| Method | Model | Input | Output resolution | Option to run without external reference | Automatic cancer cell identification | Availability (Tested version) |
| --- | --- | --- | --- | --- | --- | --- |
| InferCNV | HMM & Bayesian MM | Expression | Per gene and subclone (groups of cells) | No | No | BioConductor (v1.10.0) |
| CONICSm at | Mixture Model | Expression | Per chromosome arm and cell | Yes | No | GitHub (v0.0.0.1) |
| CaSpER | expression HMM & BAF signal shift | Expression & Genotypes | Per segment and cell | No | No | GitHub (v0.2.0) |
| copyKat | integrative Bayesian segmentation | Expression | Per gene and cell | Yes | Yes | GitHub (v1.1.0) |
| Numbat | haplotyping AFs & combined HMM | Expression & Genotypes | Per gene and subclone (groups of cells) | (Yes) | Yes | CRAN (v1.2.2) |
| SCEVAN | segmentation with variational region growing algorithm | Expression | Per segment and subclone (groups of cells) | Yes | Yes | GitHub (v1.0.1) |

The output of the CNV prediction depends on the method (**Table 1**). Half of the methods report the results per cell (CONICSmat, copyKat and CaSpER), while InferCNV, SCEVAN and Numbat group cells into subclones with the same CNV profile. Also the resolution differs, with CONICSmat reporting the results only per chromosome arm, and all other methods either per gene or per segment consisting of multiple genes. Several of the methods have two possible outputs: a discrete CNV prediction and a normalized expression score; in these cases both outputs were included separately in the evaluation and abbreviated marked with “(CNV)” and “(Expr)”, respectively. More details can be found in the **Supplementary Methods**.

We tested all scRNA-seq CNV callers on 15 different single cell RNA-seq datasets (**Figure 1**), comprising eleven cancer cell lines (nine gastric cell lines, one human breast cancer line (MCF7) and a colorectal adenocarcinoma line (COLO320)), three primary tumor samples (two basal cell carcinoma (BCC) samples and one multiple myeloma (MM) sample) and one diploid dataset (peripheral blood mononuclear cells (PBMCs)) (**Supplementary Table 1**).

**Figure 1.**
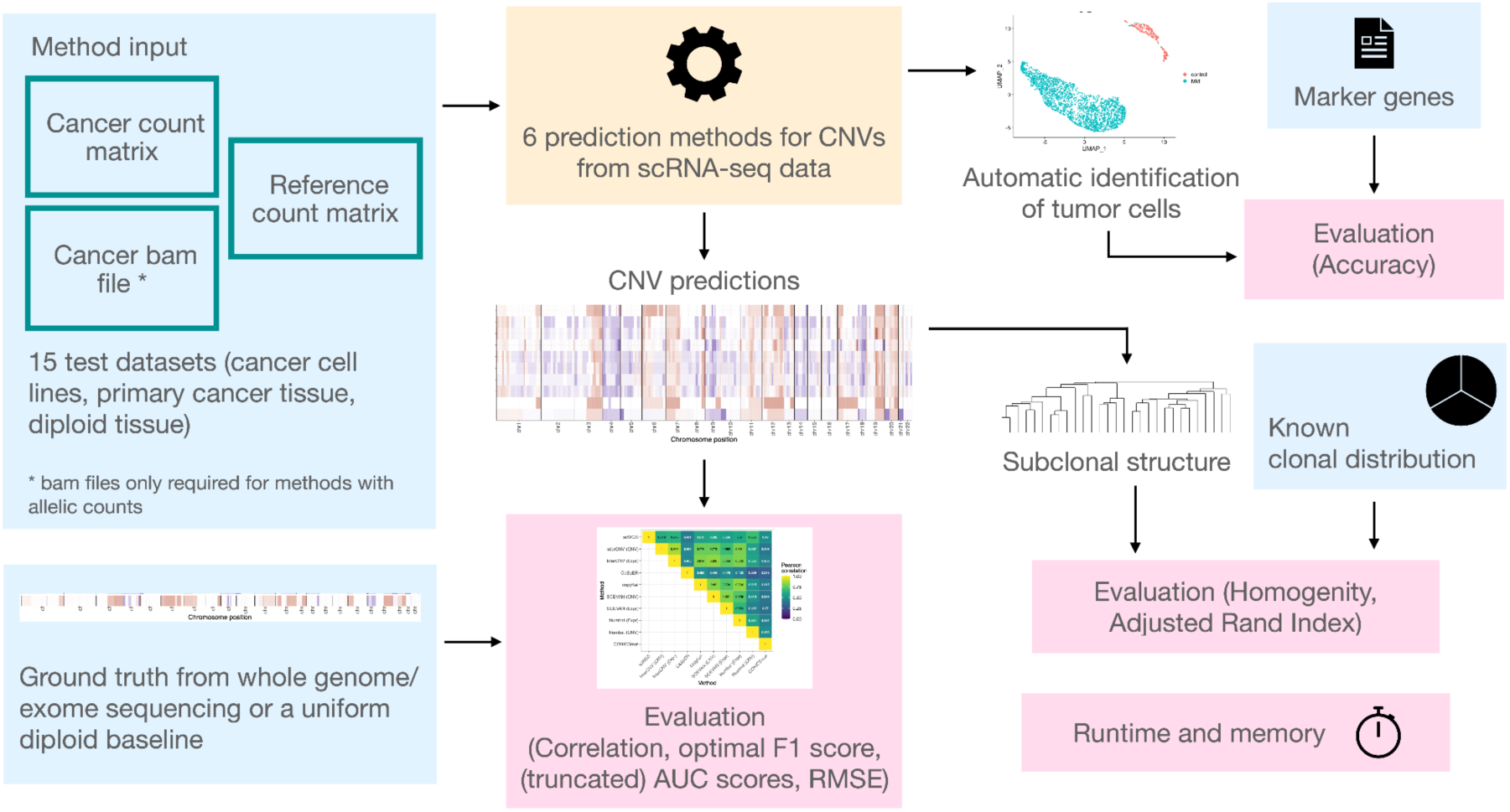
Overview schematics of the benchmarking workflow. Input data in blue, evaluation results in pink.

Different metrics were included for benchmarking, most of them based on the comparison with a ground truth for the CNVs. We obtained this ground truth from either (sc)WGS or WES data. Since the scRNA-seq methods are only able to predict the CNV status for genomic regions comprising genes, while the WGS ground truth covers (nearly) the complete genome, we could only compare modalities in gene regions. As the ground truth was not measured in the same set of cells as the scRNA-seq, and was obtained from bulk measurements in most cases, we combined the per-cell results from the scRNA-seq methods to an average CNV profile, called pseudobulk, before the comparison.

We applied threshold-independent evaluation metrics using the correlation and AUC scores. For the AUC scores, predictions were evaluated separately for gain versus all and loss versus all, resulting in two scores. Not the complete range of thresholds is biologically reasonable to classify regions as gains or losses, respectively, as every method defines a baseline score. For this reason, we chose to implement a truncated version of the AUC scores where only thresholds up to the baseline score were evaluated for losses and only scores higher than the baseline score were evaluated for gains (see **Methods**). With the same systematic, we evaluated the optimal gain and loss thresholds based on a multi-class F1 score **(Supplementary Figure 1)**, testing again only biologically meaningful gain and loss thresholds. These thresholds were then used to obtain sensitivity and specificity values for gains and losses.

Every scRNA-seq method requires a set of euploid reference cells to normalize the expression of the analyzed cells. For the primary tissue samples, the common assumption is that the measured tissues are a mixture of tumor and normal cells, of which the later can be used as reference. Some methods rely on cell type annotations provided by the user to specify the reference, while other methods provide two options, user-provided cell type annotations or automatic detection of normal cells. To ensure reproducibility between methods, cells were annotated manually into tumor and healthy cells per sample, and the same healthy cells were used as reference for all methods unless specified otherwise. For the cancer cell lines there exists no directly matched reference cells and therefore we chose, for each dataset, a matched external reference dataset with healthy cells from the same or at least very similar cell types (**Supplementary Table 1**). Since the choice of the reference euploid dataset used for normalization may affect the final CNV calling results, we tested the impact of different references on the prediction quality.

Another challenge for scRNA-seq callers is the ability to detect completely euploid datasets, with no presence of CNVs. For all methods, their performance on an euploid dataset was not evaluated in the associated publications. However, also the identification of the lack of CNVs is an important asset. For this reason, we included an euploid dataset in our performance test and calculated the mean square error deviation for every method compared to a diploid reference genome. Thereby, we explored how the performance changes depending on the choice of the reference datasets. Furthermore, for the methods with automatic detection of normal cells, we estimated the accuracy of this feature, by comparing the methods’ cell assignment to the ground truth cell type obtained from the analysis of the scRNA-seq data.

All the tested methods can detect heterogeneity in the analyzed samples. CaSpEr, CopyKat and CONICSmat estimate the CNV profiles per cell, which can be clustered afterwards into subclones, while Numbat, InferCNV and SCEVAN cluster the cells already during the analysis to improve the CNV prediction. To explore how well the methods can map cells to separate sub-clones with distinct CNV profiles, we mixed into one dataset patient data from tumors that have been shown to display high inter-individual heterogeneity in their CNV profiles. Running every method on this dataset, we quantified their ability to recover the different donors as different clones.

All evaluations were set-up within a snakemake pipeline, so that both new methods and new datasets can be easily integrated into the benchmarking.

### Benchmarking scRNA-seq CNV prediction compared to genomic ground truth

We evaluated all CNV callers on 14 different cancer datasets using various metrics. On average, Numbat (Expr), copyKat and InferCNV (Expr) had the highest maximal F1 scores (between 0.58 to 0.56), and also scored high for all other metrics (**Figure 2A, Supplementary Figure 2**). However, performance differences between most of the callers on the same dataset remained small, and all metric scores for all methods showed a large standard deviation across datasets (**Figure 2A**, **Supplementary Figure 2)**. Due to this large spread, no method can be seen as clearly superior to the others.

**Figure 2.**
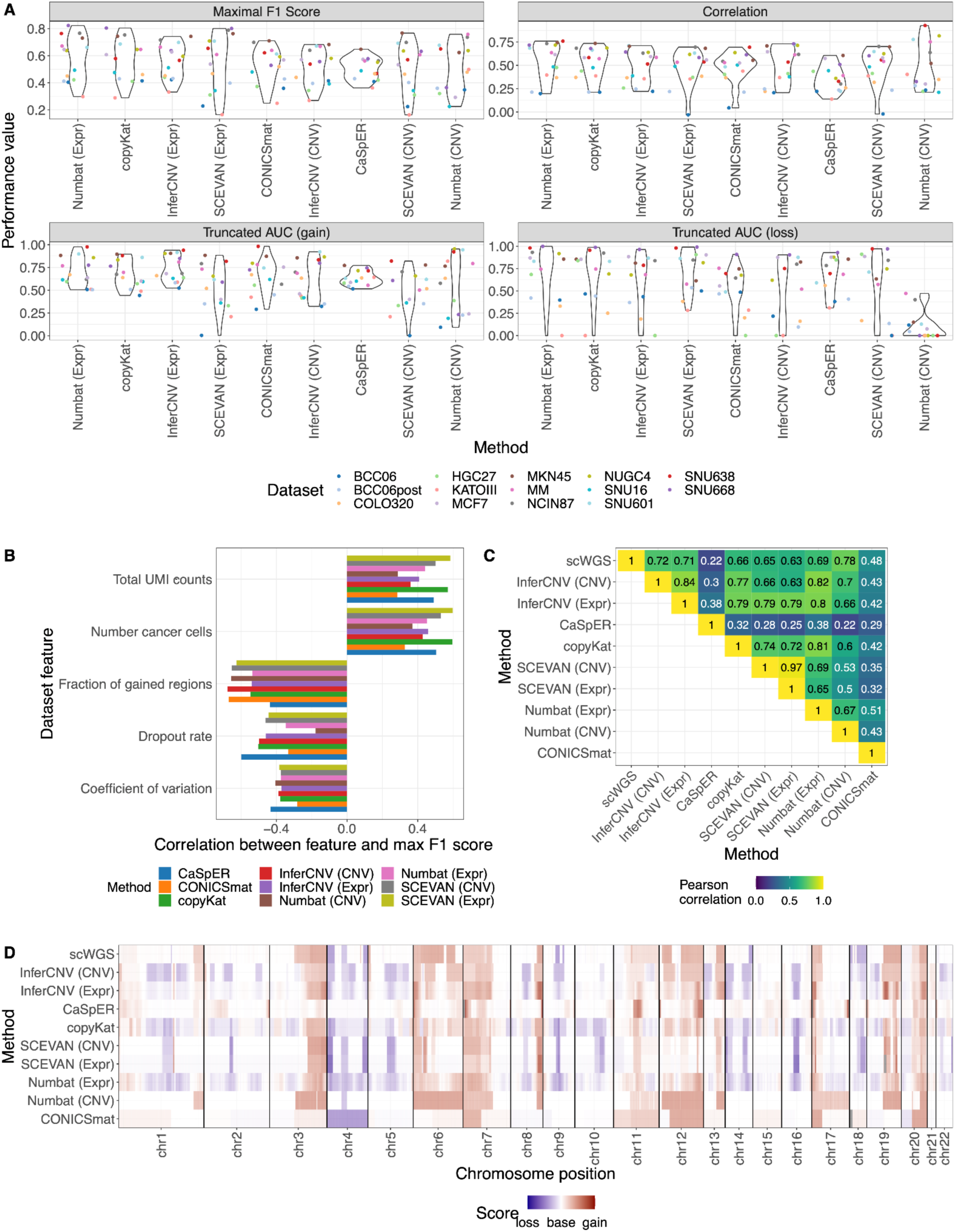
General performance evaluation on aneuploid datasets. (A) Performance comparison across all datasets. (B) Impact of dataset characteristics on the performance (maximal F1 scores). The features here show the total number of UMI counts and the number of cells in the cancer dataset, the mean dropout rate per cell, the mean coefficient of variation across genes and the fraction of gained regions from all ground truth annotated regions. Other evaluated features are shown in the supplement (**Supplementary Figure 4**). (C) Method comparison within the SNU601 dataset. (D) Karyogram of the SNU601 dataset, every method score was scaled to have the same standard deviation.

The different metrics give insights into different aspects of the prediction. The maximal F1 score puts equal weight on predicting all three CNV classes (gain, base and loss), independent of their occurrence in the dataset. This is important as most of the datasets have more gains than losses, and some of the cell lines (HGC27, KATOIII and SNU16) have very extreme profiles with more than 75% of gained regions (**Supplementary Figure 3**). A method/dataset combination with high maximal F1 score indicates that the method was able to predict all three classes accurately. The AUC and truncated AUC scores evaluate instead the separate performance between calling gains and losses. The correlation score indicates how well the overall genome-wide profile is recovered, and may reflect only the majority call. We see that generally the CNV callers are better at predicting gains (**Figure 2A, Supplementary Figure 2**). For example, in the extreme SNU638 cell line, which contains gains but nearly no losses (**Supplementary Figure 3**), Numbat (CNV) predicts the gains very well, visible in high gain sensitivity and precision (> 0.9), but it does not identify any loss regions (loss sensitivity = 0) (**Figure 2A**). For this reason, the maximal F1 score is only 0.65. However, the method exhibits a very high correlation (0.93) in that dataset, because despite not identifying any loss region at all, the overall profile was correctly predicted (**Supplementary Figure 2**).

To study the variable performance between datasets, we explored different data characteristics influencing the CNV calling (**Figure 2B, Supplementary Figure 4**). We saw a positive correlation between performance (maximal F1 score) and number of cancer cells, as well as read coverage (UMI counts). Vice versa, the performance was negatively correlated with the dropout rate, and moderately with the gene expression variation between cells. The strongest negative correlation however was observed with the fraction of genomic aberrations: the larger the fraction of the genome in the gain state, the more difficulties all methods had to infer the correct CNV profile. This is probably due to the fact that all methods seemed to have a problem identifying the baseline ploidy in these extreme cases (**Supplementary Figure 3+5**). In summary, several dataset characteristics explained the deviations in performance between datasets, and all methods were similarly affected.

For some of the CNV callers, the predictions tended to agree more between methods than when compared to the genetic ground truth (**Figure 2C+D, Supplementary Figure 5+6**). This was in particular the case for InferCNV, copyKat, SCEVAN and Numbat (Expr). This could be caused by true CNV differences between the scRNA-seq and genetic datasets, because the cells analyzed as genetic reference were not the same ones used for the scRNA-seq analysis. It could also reflect technical and biological biases of scRNA-seq data, which were picked up by all top performing methods similarly, such as problems with lowly expressed genes, or with pathways upregulated in cancer.

The methods differed in their data quality filtering steps, which ultimately lead to a different number of included genes and included cells in the CNV analysis. A more lenient gene expression filtering leads to more annotated genomic regions for CNVs. In our evaluations, we compared the CNV callers using only the overlap of all the considered regions. We therefore tested the change in performance when considering all covered regions per method, instead of only the intercept, using the SNU601 dataset, and observed no change in performance (**Supplementary Figure 7**). CaSpER and CONICSmat kept the most genes (**Supplementary Figure 8**). Despite calling CNVs for a larger portion of the genome, the permissive expression filter can negatively affect a method’s performance. This is the case for CaSpER, one of the methods with the lowest correlation values on average and no maximal F1 score above 0.65 on any dataset. To show how the expression filtering influenced performance, we exemplarily ran CaSpER with a more strict cutoff on the SNU601 cell line (keeping 8,907 genes instead of 13,196 genes) and saw a clear improvement in the CNV calling results (**Supplementary Table 2**). CONICSmat might not be as affected by the lenient expression cutoff, as it provides CNV predictions per chromosome arm, while all other methods allow for a far higher resolution. In general, an extensive parameter optimization of all methods is out of the scope of this benchmarking. We ran each method with recommended default parameters. The users should however be aware that methods might perform better with other parameters.

Additionally to the pseudobulk evaluations, we also checked the per cell estimates. For the SNU601 cell line, the per cell prediction results showed mostly minor differences between cells or subclones (**Supplementary Figure 9+10**). For all methods, the pseudobulk CNVs were closer to the ground truth compared to the per-cell results, probably due to the reduction of noise. Still, the mean per-cell correlation was over 0.45 for all methods except CaSpER and CONICSmat. So the per-cell CNV profiles of the methods are reasonable to use in downstream analysis, although an aggregation to subclonal or dataset level provides more reliable results.

### Benchmarking CNV prediction on euploid samples

Another important criterion for CNV prediction algorithms, which is usually not explored in the original methods papers, is the correct identification of euploid datasets, which display no CNVs. To consider this, we included in our benchmarking a diploid dataset consisting of CD4+ T cells from a PBMC dataset^21^, combined with four different reference datasets. Defining a matched reference dataset for the CNV prediction is a common and crucial step for all scRNA-seq CNV calling methods. It should consist of an euploid version of the same (or similar) cell type, in order to normalize the tested cells and distinguish expression changes caused by CNVs from cell type specific expression changes.

We randomly selected 50% of CD4+ T cells for the analysis, and the four reference datasets consisted of the remaining 50% of CD4+ T cells from the same sample, CD14+ Monocytes from the same sample (exemplifying the use of another cell type for normalization) (**Supplementary Figure 11A+B**); as well as CD4+ T cells and CD14+ Monocytes from another dataset^22^ (exemplifying the situation where normal cells of the same cell type are not captured in the sample) (**Supplementary Figure 11C+D**).

We calculated the Root Mean Square Error (RMSE) between the method scores and a diploid baseline (see **Methods**). When using CD4+ T cells from the same dataset as reference, all methods showed medium to good performance, visible from the karyogram plots and the low RMSE scores (**Figure 3A, Supplementary Figure 11A**). CONICSmat performed the worst with the highest divergence from a diploid genome (RMSE=0.50), followed by CaSpER (RMSE=0.47). InferCNV and copyKat found minimal CNV presence (RMSE=0.07 and 0.03 respectively), while Numbat (CNV) was the only method to identify a fully diploid genome (RMSE=0).

**Figure 3.**
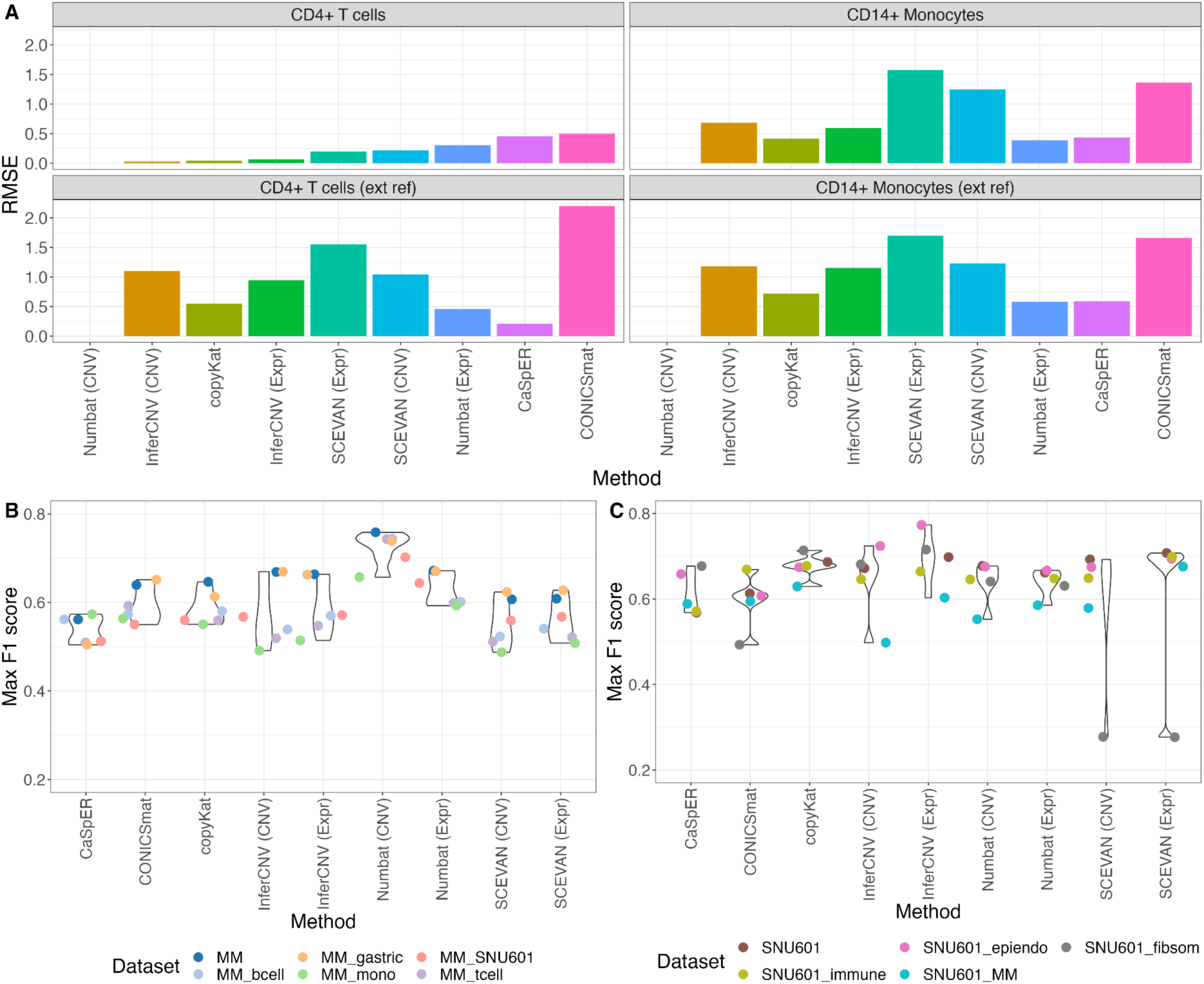
A. Root mean squared error (RMSE) between CNV predictions of each method and diploid baseline as ground truth. The methods with lowest RMSE perform best. Panel title shows chosen reference cells. **B+C**. Performance of the methods with different reference datasets for the MM dataset (B) and the SNU601 dataset (C). For the MM dataset, we tested a second healthy PBMC dataset, which we split into B cells, T cells and Monocytes (mono). We additionally tested a healthy gastric dataset and a gastric cancer dataset (SNU601). For the SNU601, we tested a second healthy gastric reference, which we split into three groups, epithelial and endothelial cells (epiendo), fibroblasts and smooth muscle cells (fibsom) and immune cells. We also tested the MM dataset as reference.

The RMSE values rose considerably when using CD14+ Monocytes instead as normalizing reference (**Figure 3A, Supplementary Figure 11B**). Here, copyKat as well as the methods that include AF information, i.e. Numbat and CaSpER, performed clearly better (RMSE<0.5). Numbat (CNV) was again the only method able to identify a fully diploid genome (RMSE=0). All other methods identified a large portion of the genome as non-diploid.

The RMSE values were much higher when using an external reference dataset^22^, independent of whether we used CD4+ T cells or CD14+ Monocytes as diploid reference. Again, Numbat, CaSpER and copyKat were the best performing methods (**Figure 3A, Supplementary Figure 11C+D**). Interestingly, Numbat (CNV) was again able to identify a fully diploid genome.

The two PBMC datasets used here were both generated with the 10X Genomics technology, however with different versions and in different laboratories, which likely had an impact on the CNV results. These technical factors cannot be overcomed by batch integration, as raw counts are required as input for calling CNVs. Even small mapping differences can influence the CNV calling performance. To show that, for the second reference PBMC dataset we repeated the analysis with the same data but mapped with an older version of CellRanger. This reduced the performance even more (**Supplementary Figure 12**).

Our results show that in general, most methods can identify diploid samples given that an appropriate reference dataset with the same cell type is provided. In cases where closely matching reference cells are not available or at least not easily identifiable, CNV calling methods with allelic information are a good option, as they are less affected by wrongly matched reference datasets. Numbat (CNV) consistently detected a fully diploid genome in all tested scenarios.

### Benchmarking the impact of the reference on CNV detection for aneuploid datasets

In real applications with cancer data, the identification of a diploid population of the same cell type is more challenging than for the euploid example. Within tumor microenvironments measured in primary samples, there exists a mixture of many cell types. Niche cells can also carry CNVs, despite not being cancer cells, which in these cases would distort CNV calling if using them as reference^23^. In the analysis of cell lines, no natural matched healthy cells exist and always an external reference is required. As discussed before, when the reference euploid dataset comes from a different sample, batch integration to minimize technical variation cannot be performed before CNV calling.

To better assess the influence of the reference dataset on the CNV results for the cancer samples, we analyzed two of the cancer datasets, MM and SNU601, with different references (**Figure 3B+C**, **Supplementary Figure 13**). For the MM primary cancer sample, the reference cells in the default analysis were the healthy cells from the tumor microenvironment. Here, additionally, we also tested different PBMC cell types as reference (T cells, B cells and monocytes from^21^), which surprisingly only lead to a moderate decrease in the quality of the CNV calls (**Figure 3B**, **Supplementary Figure 13A**). We challenged the methods further by including as reference a healthy gastric dataset^24^ (mix of fibroblasts, immune cells (B+T cells), endothelial cells, enteroendocrine cells, Chief cells, pit mucous cells, and intestinal metaplasia) as well as gastric cancer cells, which harbor CNVs on themselves (SNU601 cells^25^). For the euploid gastric cells, the quality of the CNV calls remained high, however it dropped when using aneuploid SNU601 cells as reference.

For the SNU601 dataset, we tested a second healthy gastric dataset^26^, considering three different cell groups as euploid reference (epithelial and endothelial cells, fibroblasts and smooth muscle cells, immune cells). In general, the CNV calling performance was very similar across references (**Figure 3C**, **Supplementary Figure 13B**), except for SCEVAN and CONICSmat, whose performance dropped when the gastric cancer cells were normalized using fibroblasts and smooth muscle cells. Additionally, we tested normalizing the SNU601 cells with the cancer cells from the MM dataset, to again exemplify how much the CNV results diverged when normalizing by a distant cell type that also contains CNVs itself (**Figure 3C**, **Supplementary Figure 13B**). Here, the performance dropped for all methods, although only moderately in most cases.

Overall, we see that the choice of the reference dataset has less effect on the CNV detection in aneuploid samples compared to euploid samples. However, if a cell type that itself is harboring aneuploidies is used as a reference, the CNV predictions become less reliable. Nevertheless, closely matching cell types show in most cases the best prediction results, and should therefore be the reference set of choice.

### Benchmarking automatic identification of tumor cells

In order to overcome the problem of selecting the appropriate reference dataset, CopyKat, SCEVAN, CONICSmat and Numbat can be run without the input of a reference (euploid) cell set. Importantly, in the previous sections these four methods were run with an explicit annotation of reference cells, to make the results comparable between methods. Here we explored how well these four methods worked without the reference annotation. We studied the CNV calling performance changes, as well as the correct identification of aneuploid cells. The methods use different strategies to call CNVs without the explicit annotation of reference cells. Both CopyKat and SCEVAN automatically identify putative normal cells in the dataset, which are then internally used as reference for the CNV calling. CopyKat first clusters the cells based on gene expression and defines the cluster with the minimal estimated variance in gene expression as the diploid cells. In contrast, SCEVAN uses public gene signatures of tumor, stromal and immune cells to identify highly confident normal cells and extends this annotation to similar cells via expression clustering; after CNV analysis cells are clustered again based on their predicted CNV profiles for the final annotation of non-malignant cells. In contrast to CopyKat and SCEVAN, CONICSmat does not automatically report identified diploid cells. It instead uses a two-component Gaussian Mixture Model per genomic region to identify regions with a different coverage profile as potential CNVs. Finally, Numbat uses an external reference if no internal is provided: the method contains a large gene expression reference set based on the human cell atlas and automatically matches for each cell the closest reference cell type. After running the CNV calling, the normal cells in the dataset are identified based on the aneuploidy probability, calculated from the posterior probabilities of the HMM. We tested these methods on three primary cancer datasets, MM, BCC06 and BCC06post (i.e. posttreatment) (**Supplementary Table 1**), which are supposed to be a mixture of cancer and normal cells. Their cell type annotations were taken directly from the publication for the two BCC dataset^27^ or were manually annotated based on known marker genes for the MM dataset.

We first investigated the agreement between the marker-gene-based cell type annotation and the annotation of tumor and normal cells which is provided by three of the four methods: CopyKat, SCEVAN and Numbat. In most cases, all methods showed very high concordance when predicting tumor vs normal cells **(Table 2)**. SCEVAN and Numbat reached values over 95% accuracy for all three scenarios. CopyKat performed very well on the MM and BCC06post datasets, but had problems with the BCC06 sample, as it classified about half of the manually annotated tumor cells as diploid (accuracy=44.3%). According to the manual cell type annotation, the BCC06 dataset has a very low percentage of diploid cells (∼2%) compared to the other datasets (16% and 47%). Probably for this reason, CopyKat does not work in that scenario. CopyKat identifies diploid cells as the cluster of cells with smallest gene expression variation, which can be incorrect when diploid cells do not build a large cluster. In contrast, SCEVAN includes tumor markers and Numbat CNV profiles, which both give additional evidence for the identification of normal cells.

**Table 2.**
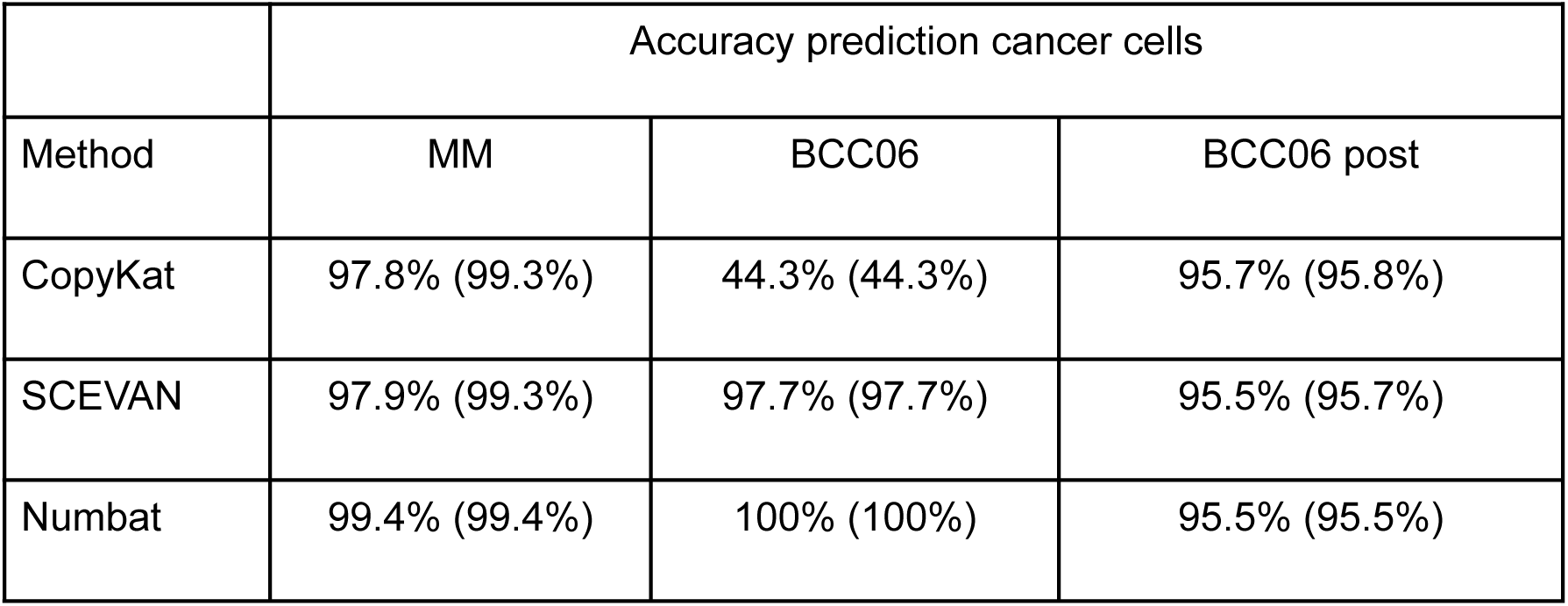
Accuracy of each method to predict the cancer cells in the respective dataset. The first number shows total accuracy including all cells, the number in brackets shows the accuracy when excluding cells defined as bad quality by the respective method.

|  | Accuracy prediction cancer cells |  |  |
| --- | --- | --- | --- |
| Method | MM | BCC06 | BCC06 post |
| CopyKat | 97.8% (99.3%) | 44.3% (44.3%) | 95.7% (95.8%) |
| SCEVAN | 97.9% (99.3%) | 97.7% (97.7%) | 95.5% (95.7%) |
| Numbat | 99.4% (99.4%) | 100% (100%) | 95.5% (95.5%) |

We then explored how much the actual CNV predictions differed when running the methods without providing the cancer cell annotation manually (**Supplementary Figure 14**). For the MM and BCC06post datasets, because the automatic identification of cancer/normal cells performed well (**Table 2**) the differences were very small between the different modes for all methods. For the BCC06 dataset however, copyKat underperformed when the reference was not provided, most likely due to the wrong identification of normal cells (**Table 2**). Surprisingly, the CNV results for SCEVAN in the BCC06 data improved when no reference was provided.

Overall, these results show that the automatic identification of reference cells is a good approach to overcome the additional effort of choosing a correct diploid reference, as the predicted results were overall very similar to the manual annotations. However, when the number of diploid cells in the sample is low, this strategy may fail, especially for CopyKat. Finally, this approach works only when normal cells are found in the dataset, which is not always the case, especially on cell lines.

### Benchmarking the identification of subclones

The big advantage of single cell data for CNV calling, compared to traditional approaches, is that subclonal structures, i.e. tumor heterogeneity, can be identified. Each of the tested methods provide information about the subclonal structure identified based on the predicted CNVs. Some methods directly define clones (Numbat, InferCNV and SCEVAN), while others produce a dendrogram (CopyKat, CONICSmat and CaSpER) that can be split to determine clones.

Obtaining independent ground truth data about the clonal structure within a cancer sample is challenging. To overcome this difficulty, we artificially created a dataset with known clonal structure by merging different cancer samples belonging to the same cancer type, but coming from different patients with different CNV profiles. To do that, we used four BCC samples^27^. CNV analysis of every patient separately shows that different patients have different CNV profiles (**Supplementary Figure 15**). Therefore, we can evaluate how effective the different methods are at sorting the patients into different CNV clusters.

All the algorithms identified several subclones in this mixed dataset (**Figure 4A**). We extracted one pseudobulk CNV profile per method and clone. The discovered pseudobulk clones clustered based on the donor rather than on the used method, suggesting that overall the methods were able to distinguish between donors.

**Figure 4.**
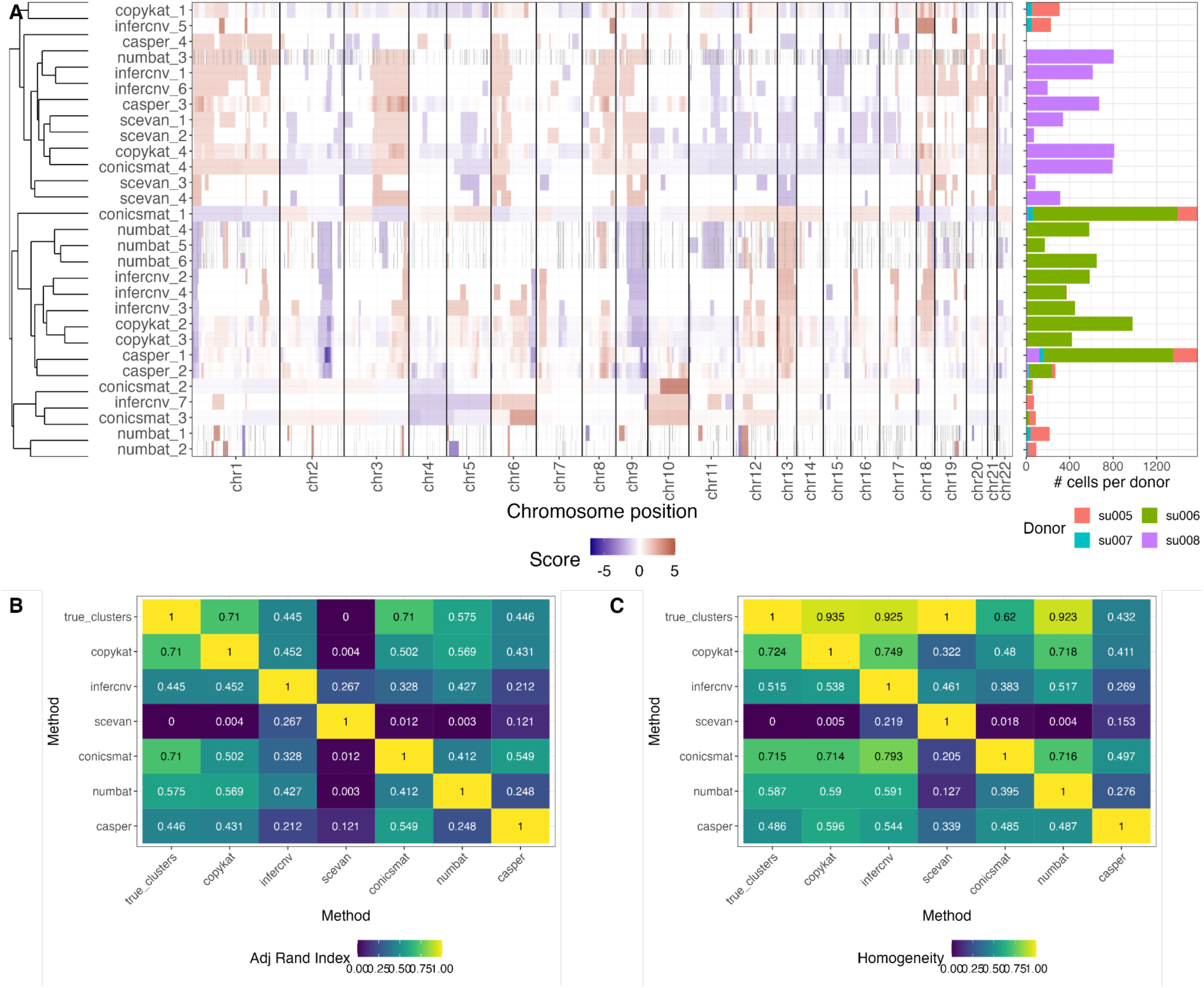
(A) Clustering of subclone CNV profiles together for all methods. Number of cells per donor visualized in the barplot on the right. (B) Adjusted Rand Index and (C) Homogeneity scores between the true clonal structure and the clonal structure identified by each method.

To quantify the performance of every method, we calculated Adjusted Rand Indices (ARI) and Homogeneity Scores for the identified clusters compared to the true donor composition. CopyKat, InferCNV, SCEVAN and Numbat all obtained Homogeneity Scores close to 1, meaning that most clones contained mostly cells from one donor alone. All four methods had lower ARI values (**Figure 4B**), probably caused by the fact that they identified multiple CNV clusters inside each donor, which is to be expected given the individual CNV results per donor (**Supplementary Figure 15**). However, SCEVAN had an ARI of 0. Despite providing a cell-type annotation file as input, SCEVAN identified only the cells from one donor (su008) as tumor cells (clusters scevan_1, 2, 3 and 4) and all the other donor cells were classified as normal (in this case no CNV profile was outputted for them). CONICSmat and CaSpER had lower Homogeneity Scores, as they grouped a large number of cells from all four donors together into one big clone (clone conicsmat_1 and casper_1). All methods had some problems distinguishing between donors su005 and su007, which could be caused by relatively similar CNV profiles of both donors (**Supplementary Figure 15**).

Overall, we see that CopyKat, InferCNV and Numbat are the best performing methods in terms of clonal structure identification, as all three were able to separate donors based on their CNV profiles.

### Overall comparison of the methods

In summary, the methods differed more in their additional functionalities than in their ability to predict CNVs (**Figure 5, Supplementary Table 3**), as the several performance metrics for the CNV predictions on the cancer datasets showed only minor differences. The characteristics of the dataset, with enough coverage and a not too extreme CNV profile, were far more relevant than the choice of the exact CNV caller. In contrast, differences between methods were visible in the analysis of euploid samples. Numbat and CaSpER were better at identifying completely diploid datasets, even when choosing a reference sample that was not closely matching the right cell type.

**Figure 5.**
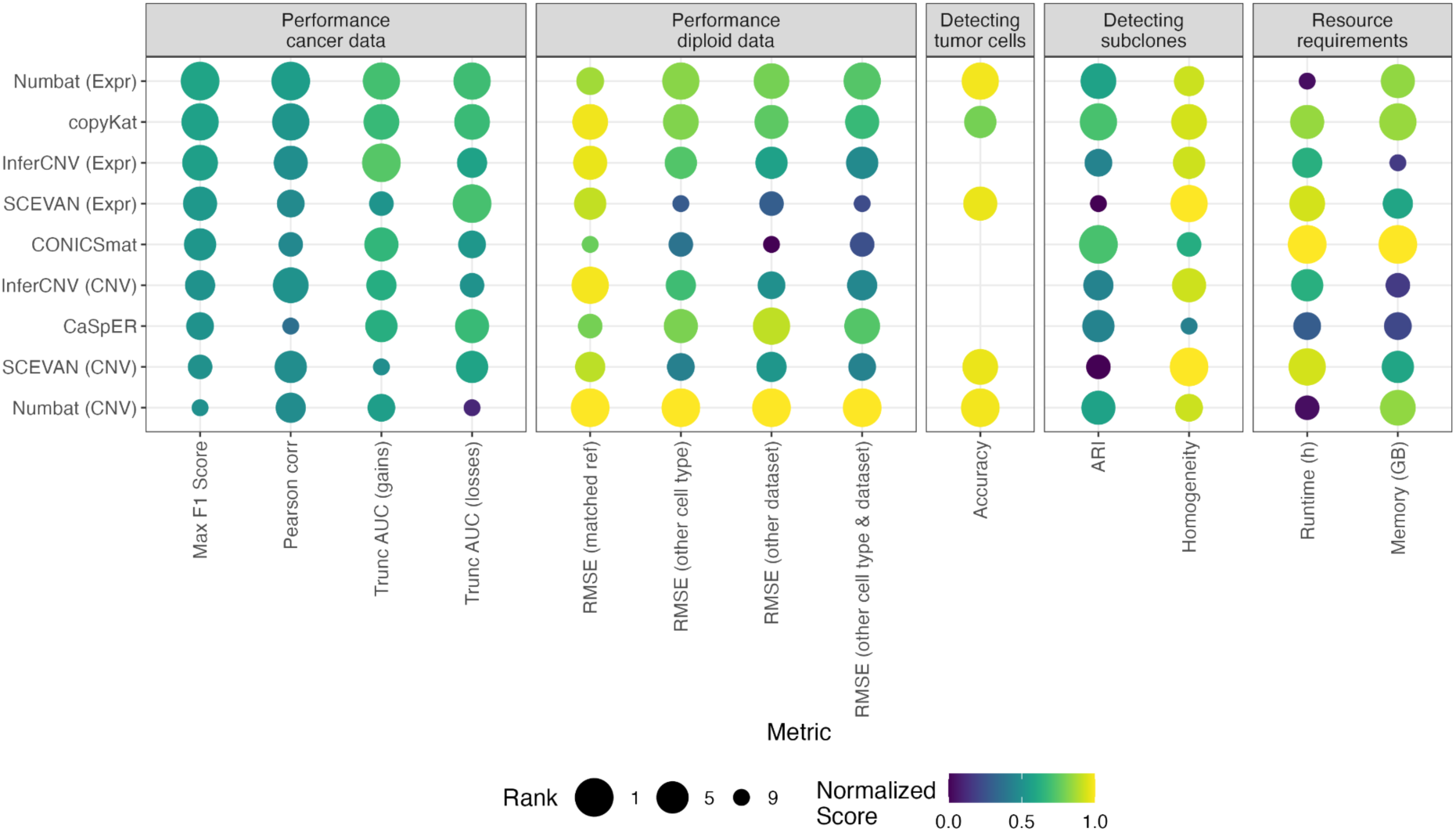
Summary of the benchmarking results. Main categories: mean CNV prediction performance for cancer datasets and diploid datasets, tumor cell classification, subclonal identification and required resources. The dot size shows the rank of each method for the respective column, with 1 being the best performing method in the category. The dot color represents a normalized score, where the values of each metric are scaled from 0 to 1, in a way that 1 is always the best value. For this, the values of RMSE, runtime and memory consumption were subtracted from one.

The automatic annotation of tumor cells in the dataset worked well for all methods which provide this feature, with best performance in Numbat and SCEVAN. Therefore, this additional functionality increases the usability of these methods. When testing the identification of CNV clones, the methods were able to assign most of the cells correctly except for CaSpER, CONICSmat and SCEVAN which did not succeed. In the case of SCEVAN, it did not distinguish any of the donors as different clones, potentially as the tested dataset was very heterogeneous.

As the performance differences were relatively small in many tested scenarios, and the dataset sizes are constantly increasing, the runtime and memory requirements of the methods should also be taken into account. We observed that the methods differed quite substantially in their resource requirements (**Figure 5, Supplementary Figure 16**). CONICSmat and CopyKat were the most efficient methods in terms of runtime and memory requirements. From these, CopyKat also showed good performance results in all categories, making it a good choice combining prediction accuracy and resource efficiency. Also SCEVAN was quite fast and memory efficient, especially given that the standard analysis already includes several downstream analyses such as gene enrichments. On the other hand, CaSpER and Numbat had the longest runtimes, as they both include allele frequency information in their analysis. Their increased robustness, shown in the detection of diploid cells, comes at the cost of more runtime. We ran each method with only one thread for better comparison, but most of the methods offer options for multi-threading, including Numbat. Finally, CaSpER and InferCNV required the most memory, which can become a problem when running datasets with many cells. As dataset sizes grow overall, the use of certain methods will become infeasible in these cases without a large computational infrastructure. Efficiently implemented methods, like CONICSmat, CopyKat and Numbat, will clearly be an advantage here.

## Discussion

This study provides a comprehensive and independent benchmarking for scRNA-seq CNV callers. We used 15 different datasets, comprising cell lines, primary cancer tissues and a diploid dataset, and extensively evaluated six methods in their ability to recover the ground truth CNV profiles, their capacity to identify the lack of CNVs in diploid samples, the correct identification of tumor vs healthy cells, and the accurate identification of subclonal CNV structure in the data. We carefully selected and designed a set of metrics, as the CNV prediction is a three-class problem with continuous scores, but a restricted biological meaningful value range. We applied either threshold independent evaluation metrics, such as the correlation and (truncated) AUC scores, or chose the thresholds which produced the maximal F1 scores.

The benchmarking did not identify one single method which outperformed all others in all tasks. While Numbat (Expr), copyKat and InferCNV (Expr) outperformed at predicting CNVs in aneuploid datasets, Numbat (CNV) outperformed all other methods at detecting the right CNV profile in euploid samples regardless of the reference diploid dataset used. The adequateness of the cell type used as diploid reference had a strong effect in the correct identification of diploid cells for all other methods. Furthermore, using cells from an external reference reduced the performance, even when using a matching cell type. This is to be kept in mind when designing experiments, if the reference euploid cells are measured separately from the aneuploid ones. The effect of the best matching reference sample was milder on the CNV discovery in aneuploid samples instead.

Dataset quality had far more impact on the CNV results than the chosen algorithmic solution. In particular, both the number of cancer cells and the total UMI counts in the dataset positively influenced the performance of all methods; while the dropout rate, the amount of gene expression variation between cells, and the fraction of the genome in the gain state negatively influenced it. In general, losses were more difficult to identify compared to gains. Specifically for the sparse single cell data, differentiation between dropouts and low coverage due to copy number losses is challenging. Furthermore, previous studies about CNVs reported that losses occur more frequently in genomic regions of low gene density^28^, another disadvantage for scRNA-seq callers.

The methods had very different performances at their other functionalities. Three of the evaluated methods, copyKat, SCEVAN and Numbat, are able to identify tumor vs healthy cells. When the number of normal cells in the sample was large, all methods excelled at their identification. However, when only few normal cells were present copyKat wrongly identified them. Finally, all methods provide information about the CNV clones present in the analyzed sample. CopyKat, InferCNV and Numbat were able to identify the right clonality, while on the opposite side of the spectrum SCEVAN was not able to disentangle any clones.

In terms of resources, CONICSmat was the most resource-efficient method (both runtime and memory), but its CNV detection performance showed weaknesses and was highly influenced by the choice of reference cells. Moreover, the method could not correctly disentangle the clones present in a mixed dataset. CopKat and SCEVAN were fast methods. CopyKat’s performance was high in all tests, except at the automatic identification of healthy cells if these were not numerous in the sample. SCEVAN instead had variable performance at the CNV identification tests and could not identify clones in a mixed sample, but it excelled at the identification of healthy vs tumor cells. InferCNV used the most memory of all tested methods but performed relatively well in all evaluated categories. CaSpER and Numbat were the most runtime-intensive methods, probably due to the incorporation of allele frequency information. Between the two, Numbat offers more additional functionalities and was overall scoring better than CaSpER in all of our tests.

We focused in our study on the basic distinction between gain, base and loss CNV classes, to homogenize results between methods. However, some methods allow more detailed differentiation of the CNV states. For example, InferCNV reports the exact ploidy up to three gained copies. Both Numbat and CaSpER report additionally the copy-neutral loss of heterozygosity, which is possible only as they include allele frequency information. Depending on the exact use case, these additional functionalities might be helpful. Methods including allele frequency information cannot be applied on data from mouse strains, which carry too little heterozygous variation.

We included in the benchmarking both primary cancer samples and cell lines, which differ slightly in their analysis, as cell lines require external reference datasets. This issue applies not only to the commercial cell lines we used in our test cases, but also to any newly established cell line in the laboratory. This makes our results, specifically for analyzing cell lines, relevant for different scenarios.

The usability of the tools was not evaluated, as an objective quantitative metric for this is difficult to obtain. All analyzed methods provided a tutorial with installation instructions and an example analysis. CopyKat, Numbat and SCEVAN can only be run for a predefined set of gene annotations, therefore they can only be applied to human and mouse datasets on a limited set of genome versions. Some of the methods, i.e. Numbat, SCEVAN and InferCNV, report discrete CNV classes; all other methods output a continuous CNV score, which is not straight-forward to discretize and interpret by the users. Moreover, methods differ largely on their downstream functions, which were not benchmarked here. As several of the methods were developed recently, an active user community can potentially help with their feedback to overcome current limitations. We tested but did not include two additional CNV calling methods, HoneyBadger^10^ and sciCNV^29^. HoneyBadger did not terminate running on the SNU601 dataset (5,880 cells) after multiple days, likely because it was not developed for droplet-based datasets with many cells. For sciCNV, there were errors in the code, which did not allow us to execute the method’s tutorial.

There is a general debate whether CNV profiles can be accurately inferred from scRNA-seq data, as the gene expression is only indirectly associated with the CNV profile and additionally influenced by other regulatory factors, and RNA-seq data covers only a small part of the genome. Nevertheless, as it is the most abundant single cell technology, prediction of CNVs from it represents an appealing possibility. In our benchmarking, we showed that all tested RNA CNV callers were able to infer the CNV profile with relative accuracy, given that the analyzed dataset had sufficient coverage and a not too extreme CNV profile.

We have previously shown that the prediction of CNVs from other omics layers, such as scATAC-seq, performs better in comparison to scRNA-seq^16^. The establishment of multi-omics methods offers promising new opportunities to improve CNV calling in the future^30^. Furthermore, CNV calling methods are currently being extended to spatial transcriptomics data^31,32^. Our benchmarking does not only provide a guideline for the analysis of scRNA-seq CNVs, but it also represents the basis to develop and improve CNV calling methods. To support the community, we have provided a reproducible Snakemake pipeline, such that users can test their datasets against all implemented methods, and developers can test the performance of their new methods against a wide range of datasets and functionalities.

## Methods

### Running the CNV RNA-seq methods

We ran each method with their default parameters or the parameters suggested for droplet-based dataset in their tutorials. Further information on each method can be found in the **Supplement Methods**. Runtime and memory consumption of each method was automatically tracked within the snakemake workflow^33^, which was used to apply each method on all datasets.

### Preprocessing the single cell datasets

The required input files for our pipeline are a count matrix with cell type annotations, which defines most importantly the cancer cells, and a bam file to estimate allele frequencies for Numbat and CaSpER. Bam files and count matrices were generated from the fastq files by running CellRanger (version 7.0.0), except for the BCC dataset, where the annotated count matrix was taken directly from GEO. The cell type annotation is only relevant for the primary cancer samples, as the cell lines are assumed to be a homogenous set of cells and all labeled as the cell line. For the BCC dataset, the published cell type annotation was used. For the MM dataset, a manual annotation was performed, using Louvain clustering and known marker genes for MM (CD38, IGHM, MZB1).

In contrast, the cell lines required all an external reference dataset. We selected them to match the original tissue of the cell line, i.e. intestine for SNU601, colon for COLO320 and breast for MCF7 (see **Supplementary Table 1**).

### Preprocessing the genomic ground truth

The raw reads of the low-pass WGS MCF7 cell line^34^, as well as the WES reads of the BCC SU006 (pre- and posttreatment)^27^ and the Multiple Myeloma (sample MM199)^29^, were first trimmed in order to remove low quality reads, using Trimmomatic (removing the first and last bases with quality less than 5, sliding window of size 4 and cumulative quality of 15, removing reads with length less than 36 bases). The trimmed reads were aligned to the human genome, version hg38, using BWA^35^. The aligned reads were filtered for duplicate reads (PCR artifacts), using the MarkDuplicates software from the Picard tools^36^. For all the WES tumor-control paired sample, CNVs were called using the GATK4^37^ Best Practices recommendations^38,39^, tuned for WES datasets. For the low-pass WGS MCF7 sample, CNVs were identified using ichorCNA^40^ and in accordance with the tool manual.

The copy number variations of the COLO320 WGS dataset^41^ was performed using CNVkit^42^, with parameters set for WGS as suggested in the manual. For the gastric cell lines, the scWGS^25^ groundtruth copy numbers were called with the same process as in ^16^ for the SNU601 cell line.

### Evaluation metrics for comparison to genomic ground truth

Different approaches were implemented to compare the continuous CNV estimates from the CNV scRNA-seq methods with the genomic ground truth datasets. Before the evaluation, the CNV scores from each scRNA-seq method are aggregated to pseudobulk scores, combining all cells annotated as tumor cells. In case the genomic ground truth was also generated from single cell data, also here the pseudobulk is calculated. Genomic CNV results were obtained in 100kB bins. The CNV scores from the scRNA-seq methods exist in different formats, either per gene or for longer segments. Each version was mapped to the 100kB bins. If multiple genes or segments overlapped with one 100kB bin, the average score was taken. This resulted in two vectors of continuous CNV scores per bin, one for the genomic ground truth and one for the CNV estimates.

We chose several threshold independent metrics for the benchmarking, namely Pearson correlation between scRNA-seq CNV calls and the genomic ground truth, and different versions of the AUC. For the AUC scores, an AUC (gain) was calculated for predicting gain versus the other CNV types (base or loss) and an AUC (loss) for predicting loss versus the other CNV types (gain or base).

However, not all thresholds are reasonable for gain or loss prediction, respectively. Every method has a baseline value, usually 0, and only loss values lower than the baseline and gain values higher than the baseline are biologically reasonable. Therefore, we calculated a truncated version of the AUCs, where we only tested thresholds that are biologically meaningful. We reported the truncated AUC loss and gain values on top of the standard AUC values.

For the AUC calculations, a discrete ground truth is required. For the bulk data, this is given directly, for the scWGS, loss values are defined by at least 50% of the cells having a loss at this segment and gain values as at least 50% of cells having a gain there.

Furthermore, to obtain sensitivity and specificity values for gains and losses, we evaluated the optimal gain and loss thresholds, respectively. For each method, the multi-class F1 score was calculated for each possible threshold combination, again limiting the search to biological meaningful gain and loss values. The multi-class F1 threshold was estimated based on the mean of the gain F1 score, the base F1 score and the loss F1 score. By giving each CNV the same weight, also the identification of rare CNV classes stays important.

### Exploring performance differences across datasets

Each tested cancer dataset was described with different characteristics. Several values were taken from the scRNA-seq count matrices: the number of cells in the dataset (*num_cancer_cells*), the mean number of UMI per cell (*mean_umi_cancer*), the total number of UMI in the dataset (*total_cancer_umi*), the mean number of non-zero genes per cell (*mean_nonZeroGenes_cell*), the mean dropout rate per cell (*mean_drop_cell*), the mean coefficient of variation across genes (*mean_coef_var*). The coefficient of variation is calculated based on the standard deviation of the log-normalized counts with the formula^43^

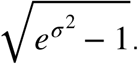

Specifically for the evaluation of Numbat and Casper, also the identification of SNPs in each dataset was explored: the total number of identified SNPs by CaSpER (*casper_nsnps*) and Numbat, respectively (*numbat_nsnps*) as well as the number of cells with at least one SNP according to Numbat (*numbat_cell_wsnp*). CaSpER identifies the SNPs only per sample, not separately in each cell.

From the genomic ground truth data, we extracted several characteristics to describe the expected CNV distribution: the fraction of loss bins and gain bins from all annotated bins (*fraction_loss*, *fraction_gain*), the fraction of CNV bins of any type, i.e. loss or gain (*fraction_cnvs*), and the number of breakpoints, where the CNV status changes (*num_breakpoints*). A higher number of breakpoints shows that there are more CNV (and potentially shorter) CNV segments. Lastly, we included the total number of genomic bins, each of size 100kB, which were evaluated for each dataset (*num_genomic_regions*).

We evaluated the impact of all dataset characteristics on the performance by calculating the Pearson correlation between each feature and the maximal F1 score.

### Evaluation of diploid samples

We evaluated how well the CNV prediction methods perform on diploid datasets by testing them on CD4+ T cells from a PBMC dataset of a healthy donor^21^. First, we annotated the cell types in the dataset using classical marker genes, following the muon tutorial^44^. Then, we randomly split the CD4+ T cells and used 50% as test dataset for the prediction in combination with four different reference datasets. As references, we tested the other 50% of the CD4+ T cells and the CD14+ Monocytes from the same dataset as well as CD4+ T cells and CD14+ Monocytes from a second PBMC dataset^22^. As the second dataset was originally mapped with an old version of CellRanger (v1.0), which did not count intronic reads, we remapped it with a newer version of CellRanger (v7.0). We additionally tested the differences when using the newly mapped matrix and the original published matrix from GEO.

For the evaluation, we assume a completely diploid ground truth, i.e. every bin is base. This requires different evaluation metrics as for the cancer datasets. We calculated here the root mean square error (RMSE) of each method from the zero baseline. As the magnitude of the scores is different, we first normalized the scores based on their standard deviation in the SNU601 dataset. This way, we can interpret the deviation dependent on a typical deviation seen in a cancer dataset.

### Performance of aneuploid datasets with different reference sets

We tested for two of the cancer datasets different reference datasets, SNU601 and MM. For each reference dataset, we repeated the whole evaluation workflow, i.e. running every method and comparing the results to the genome ground truth. For the MM dataset, we tested different immune blood cells as reference, taking the diploid dataset we evaluated before^21^. We tested three different subsets, T cells, B cells and Monocytes. Additionally, we tested two extreme references, the SNU601 dataset^25^ and the gastric reference of the SNU601^24^.

For SNU601, we tested a second gastric dataset^26^, which we splitted into three subsets dependent on the published cell type annotation. The first subset contains epithelial and endothelial cells, the second subset fibroblasts and smooth muscle cells and the third dataset all types of immune cells. Additionally, we tested the SNU601 taking the cancer cells from the MM dataset as a reference, an extreme case where the reference contains itself CNVs.

### Evaluation of automatic cancer cell identification

There are four scRNA-seq CNV calling methods which can be run without providing an explicit cell type annotation: CONICSmat, CopyKat, Numbat and SCEVAN. In the previous analyses, they were always run with a defined reference annotation to increase the comparability with the other methods. Here, each was tested a second time without explicit reference cells for the three primary tumor datasets (MM, BCC06 and BCC06post). These three datasets comprise a mixture of tumor and euploid cells. The performance was evaluated with the same metrics as in the previous analyses and the differences with and without the explicit reference cells were compared.

Three of the methods (CopyKat, Numbat and SCEVAN) additionally annotate the cells as cancer cells vs euploid cells, when running without an explicit annotation. We compared these annotations to our own manual cell type annotation for the MM dataset and to the published cell type annotations for the two BCC datasets.

### Evaluation of the identified subclonal structure

A known ground truth was required to evaluate how well each method can distinguish different CNV clones. For this, the pre-treatment data of four donors of the BCC dataset were chosen, selecting the donors with at least 20 cancer cells. First, we ran each donor separately with copyKat and clustered the per cell CNV predictions afterward to verify that the donors have indeed distinguishable CNV profiles, i.e. can be seen as separate clones.

Then a combined count matrix and bam file was generated to mix the four samples, excluding all donor information in the analysis. Afterward, the standard CNV calling pipeline was run. Numbat, InferCNV and SCEVAN directly provide a classification of cells into distinct groups, i.e. subclones, while CopyKat, CONICSmat and CaSpER return only a hierarchical clustering. For the last three methods, we cutted the tree at a height of 4 to obtain distinct groups for the evaluation.

We used two different metrics to compare the clusters to the ground truth of patient annotations, the adjusted Rand Index (ARI) and the Homogeneity score. The ARI evaluates the similarity between both clustering results. However, some of the methods split the cells of one patient into multiple clusters, so potentially multiple subclones per patient, which we can not verify with this dataset. This problem is overcome with the Homogeneity score, which evaluates whether all cells from one cluster belong to the same group, here to the same patient. Splitting one patient into several subclones is thereby not punished.

## Data availability

For the nine gastric cell lines^25^, the scWGS data was downloaded from the SRA repository, with accession number PRJNA498809. The corresponding scRNA-seq data was downloaded from the SRA repository RRJNA598203 and the control samples for the scRNA-seq^24^ data from the GEO repository with accession number GSE150290. A second gastric dataset^26^ was tested as an alternative reference for the SNU601, downloaded from GEO (accession number GSE159929, the stomach dataset GSM4850590).

For the breast cancer cell line MCF7^34^, ultra low pass whole genome sequencing data was downloaded from the ENA, project PRJNA398960 (Biosample SAMN07519582), while the scRNA-seq dataset was download for GEO (accession number GSM3142233). As a reference for the MCF7 cell line, the mammary gland dataset from the Tabula Sapiens cell a t l a s ^4^ ^5^ was used (https://figshare.com/articles/dataset/Tabula_Sapiens_release_1_0/14267219).

The multiple myeloma (MM) whole exome sequencing dataset^29^ was downloaded from the GEO (accession number GSM4200481 for the control sample and GSM4200480 for the tumor sample), while the corresponding scRNA-seq dataset was downloaded from GEO (accession number GSM4200471).

The COLO320HSR whole genome sequencing dataset^41^ was downloaded from the SRA accession PRJNA506071 (sample SRS4831935), the multiome data^46^ of the same cell line from the SRA accession PRJNA672109 (sample SRS7587918), and the control scRNA-seq samples^47^ were acquired from https://www.gutcellatlas.org/.

The basal cell carcinoma (BCC)^27^ scRNA-seq samples were downloaded from the GEO repository with number GSE129785 (GSM3722057 for su006 and GSM3722057 for su006 post, GSM3722053 for su005, GSM3722058 for su007 and GSM3722063 for su008). For the BCC sample su006 WES data were downloaded from the SRA accession PRJNA533341 (sample SRS4645190).

Lastly, two PBMC datasets that were tested as diploid samples for CNV calling, were downloaded from 10x Genomics (https://www.10xgenomics.com/datasets/pbmc-from-a-healthy-donor-no-cell-sorting-10-k-1-standard-2-0-0) and GEO accession GSE96583^22^ (sample GSM2560248, batch A).

## Code availability

All code written for this study, especially the benchmarking pipeline with snakemake, can be found on github https://github.com/colomemaria/benchmark_scrnaseq_cnv_callers

## Acknowledgements

We thank Dr. Gregor Miller and Ronan Le Gleut from the Core Facility Statistical Consulting at Helmholtz Munich for statistical advice. We thank the BMC Bioinformatics Core Facility for providing access to their HPC cluster.

## Author contributions

MCT designed the study. KTS, AS and MLR prepared the data. KTS and DH implemented the benchmarking pipeline. KTS and MCT wrote the manuscript with support from AS and MLR. All authors reviewed the final manuscript.

