## Supplementary Material for "Benchmarking scRNA-seq copy number variation callers"

### Supplementary Material - Benchmarking scRNA-seq copy number variation callers

Katharina T. Schmid [1], Aikaterini Symeonidi [1,2], Dmytro Hlushchenko [1], Maria L. Richter [1], Maria Colomé-Tatché [1,2]
1 Biomedical Center (BMC), Physiological Chemistry, Faculty of Medicine, LMU Munich, Planegg-Martinsried, Germany
2 Institute of Computational Biology, Computational Health Center, Helmholtz Zentrum München, German Research Center for Environmental Health, Neuherberg, Germany

#### Supplementary Methods

##### InferCNV

InferCNV identifies CNVs based on an HMM model, which is followed by a Bayesian Mixture Model to remove potential false positive predictions. It is available as an R-package on BioConductor, we evaluated version 1.10.0. We ran the default 6-state HMM model with the analysis mode "subclusters" to identify potential different CNV states between subclones. The default analysis mode is "sample", but the documentation states that the subcluster mode is slower, but more accurate, so we chose this. Gene annotations were taken from the tutorial website, based on Gencode hg38 annotations. The filtering parameter "cutoff" was set to 0.1, which represents the minimal average count threshold per gene among the reference cells. This value was recommended for 10X data in the tutorial. All other parameters were kept to their default values.
The final results comprise two different outcomes, a matrix with relative expression intensities and a matrix with pruned CNV predictions in 6 states. Both outputs are evaluated independently as InferCNV (Expr) and InferCNV (CNV), respectively. The 6 outcome states are grouped into gain - base - loss with states 1 and 2 being loss, 3 being base and 4 to 6 being gain.

##### CONICSmat

CONICSmat is an adaptation of the tool CONICS, which uses DNA sequencing data in combination with scRNA-seq data to analyze CNVs. In contrast to CONICS, CONICSmat requires no genomic data. Therefore, we included only CONICSmat in our analysis, as the inputs are comparable to the other tools.
CNVs are estimated based on a two component Gaussian Mixture Model (GMM) of the expression data of each cell and region. The regions can either be predefined as potential copy number alterations based on genomic data or, as in our case, without any additional data, each chromosome arm is defined as one region. CONICSmat requires a dataset with euploid and aneuploid cells to fit the two component GMMs, but an annotation of euploid cells is optional. The tool can also be run without providing an explicit reference. We tested both options, once with and once without defining an explicit reference. Regions where the two component GMM is statistically more likely than a uniform distribution are defined as CNV regions and their type (gain or loss) is assigned based on the expression levels. We tested the R package version v0.0.0.1 in our benchmarking.

##### CaSpER

CaSpER combines the expression signal with loss of heterozygosity information, based on SNPs called from the single cell data. It performs multi-scale smoothing of the expression signal followed by a 5-state HMM model for each scale. The outcome is combined with a multi-scale smoothed BAF shift signal to confirm the intermediate HMM CNV states. CNV calls are obtained for each combination of expression HMM and BAF shift signal. We apply 3 different scales for both expression and BAF signal, resulting in 9 CNV calls for each segment. Following the tutorial, summary gains and losses are defined, when they are called at least 7 of 9 possible times.

Preprocessing of the datasets was done with Seurat analogously to the tutorial; the cells were normalized to 1 million counts per cell and logarithmized before running CaSpER. We chose an expression threshold of 0.1 (parameter name “expr.cutoff”), as applied in the 10X specific tutorial of CaSpER. We tested version 0.2.0 of the R package.

As an additional remark, the complete CaSpER object, generated by the package, can become very large. To save memory, we stopped storing this object as an RDS file in the end.

##### copyKat

CopyKat attempts to overcome the issue of manually defining reference cells by automatic annotation of normal cells in mixed samples from tumor microenvironments. Normal cells are labeled based on the assumption that they cluster together and display the lowest variance compared to tumor cells. The CNV profile is identified via an integrative Bayesian segmentation approach.

We tested the version 1.0.1, using the default parameters for filtering and segmentation. The gene position file is provided internally, but only for human (hg20) and mouse (mm10). The method was evaluated twice, once with specified reference cells and once without, so that the reference cells were identified automatically. The output file contains relative copy number ratios (instead of discrete CNV states).

##### Numbat

Numbat aims to improve CNV detection by combining expression and genotype information. In contrast to CaSpER, it uses population-based phasing to get haplotype information. THis is more sensitive compared to using SNPs directly, as it can overcome the sparsity to a certain extent. Expression and haplotype frequencies are combined in an HMM model. The method also infers the phylogenetic tree during the CNV prediction, so that information from cells belonging to the same subclone can be aggregated to pseudobulk, again to reduce the sparsity. For this, it alternates between CNV prediction and reconstructing the phylogenetic tree based on the identified CNVs. In the end, tumor and normal cells can be classified based on posterior probabilities for the baseline CNV.

Numbat provides a large external reference dataset for the analysis with many cell types from the human cell atlas. However, we generated our own reference set per dataset, so that the same reference is used between all methods and the comparability is increased. We tested the R package version 1.2.2. To check how much the allele frequency information adds to the CNV detection, we included the normalized expression profiles of the single cells as Numbat (Expr), which does not contain any genotype information, and the clone-level pseudobulk CNV profiles as Numbat (CNV), which contains both expression and genotype information.

##### SCEVAN

Similar to CopyKat, SCEVAN offers the option to automatically identify normal cells as a reference, however with a different approach. A set of so-called confidential non-malignent cells is identified based on gene profiles of malignant and non-malignant cells and then this set is extended to similar cells based on expression clustering. With this reference, genes

are normalized and smoothed before a joint segmentation across all cells together identifies common breakpoints. Cells are clustered afterwards to identify both healthy diploid cells and then different subclones. Afterwards, one CNV profile is obtained per subclone.
SCEVAN includes in its workflow several further analyses such as pathway enrichment analysis, which can facilitate the interpretation of CNV results, but are not included in our benchmarking. We tested version 1.0.1 of the R package. We evaluated the normalized expression per cell, aggregated to pseudobulk, in SCEVAN (Expr) and the final CNV profile per subclone, again aggregated to pseudobulk, in SCEVAN (CNV). The final CNV prediction was simplified to 0 and 1 both being loss, 2 being base and 3 and 4 being gain. The method was evaluated twice, once with specified reference cells and once without, so that the reference cells were identified automatically.

#### Supplementary Tables

| Dataset | Reference | Sample Type | Cancer Type | Technology | Reference samples | Ground Truth | Number cells (cancer + ref) |
|---|---|---|---|---|---|---|---|
| COLO320 | Hung et al, 2021 | Cell line | Colon adenocarcinoma | 10x multiome sequencing | Healthy human adult from gut cell atlas (Elmentait et al., 2021) | WGS | 4372 + 5000 |
| HGC27 | Andor et al., 2020 | Cell line | Gastric carcinoma | 10x scRNA-seq | 5 control samples from a gastric cancer study (Kim et al., 2022) | scWGS | 1728 + 4813 |
| KATOIII | Andor et al., 2020 | Cell line | Gastric adenocarcinoma | 10x scRNA-seq | 5 control samples from a gastric cancer study (Kim et al., 2022) | scWGS | 3637 + 4813 |
| MCF7 | Ben-David et al., 2018 | Cell line | Breast cancer | 10x scRNA-seq | Mammary gland data from Tabula Sapiens (The Tabula Sapiens Consortium et al, 2022) | WGS | 1726 + 5630 |
| MKN45 | Andor et al., 2020 | Cell line | Gastric adenocarcinoma | 10x scRNA-seq | 5 control samples from a gastric cancer study (Kim et al., 2022) | scWGS | 4221 + 4813 |
| NCIN87 | Andor et al., 2020 | Cell line | Gastric adenocarcinoma | 10x scRNA-seq | 5 control samples from a gastric cancer study (Kim et al., 2022) | scWGS | 4753 + 4813 |
| NUGC4 | Andor et al., 2020 | Cell line | Gastric adenocarcinoma | 10x scRNA-seq | 5 control samples from a gastric cancer study (Kim et al., 2022) | scWGS | 4991 + 4813 |
| SNU16 | Andor et al., 2020 | Cell line | Gastric adenocarcinoma | 10x scRNA-seq | 5 control samples from a gastric cancer study (Kim et al., 2022) | scWGS | 1444 + 4813 |
| SNU601 | Andor et al., 2020 | Cell line | Gastric adenocarcinoma | 10x scRNA-seq | 5 control samples from a gastric cancer study (Kim et al., 2022) | scWGS | 5880 + 4813 |
| SNU638 | Andor et al., 2020 | Cell line | Gastric adenocarcinoma | 10x scRNA-seq | 5 control samples from a gastric cancer study (Kim et al., 2022) | scWGS | 1117 + 4813 |
| SNU668 | Andor et al., 2020 | Cell line | Gastric adenocarcinoma | 10x scRNA-seq | 5 control samples from a gastric cancer study (Kim et al., 2022) | scWGS | 7005 + 4813 |
| MM | Mahdipour-Shirayeh et al., 2022 | Primary cancer sample | Multiple myeloma | 10x scRNA-seq | Normal cells within the dataset | WES | 2003 + 257 |

| | | | | | | | |
|---|---|---|---|---|---|---|---|
| BCC06 | Yost et al., 2019 | Primary cancer sample | Basal cell carcinoma | 10x scRNA-seq | Normal cells within the dataset | WES | 1403 + 33 |
| BCC06post | Yost et al., 2019 | Primary cancer sample | Basal cell carcinoma | 10x scRNA-seq | Normal cells within the dataset | WES | 320 + 280 |
| PBMC (CD4 T cells) | 10X Genomics | Diploid | | 10x scRNA-seq | Subsample of CD4T cells / Monocytes from the same dataset | Diploid baseline | 535 + 535/1842 |

**Supplementary Table 1.** Evaluated datasets

| Expression threshold | Pearson correlation | Maximal F1 score | Truncated AUC (gain) | Truncated AUC (loss) |
|---|---|---|---|---|
| 0.1 | 0.21 | 0.55 | 0.62 | 0.59 |
| 4.5 | 0.64 | 0.68 | 0.83 | 0.86 |

**Supplementary Table 2.** Performance differences for CaSpER depending on the chosen expression threshold for the SNU601 cell line. Recommended cutoff based on the 10X tutorial of CaSpER is 0.1, default parameter for bulk and plate-based single cell data is 4.5.

**Supplementary Table 3.** Mean performance for each method and analysis category across all respectively tested datasets - see separate xls file (SupplementaryTable3_metrics_means.xlsx)

#### Supplementary Figures

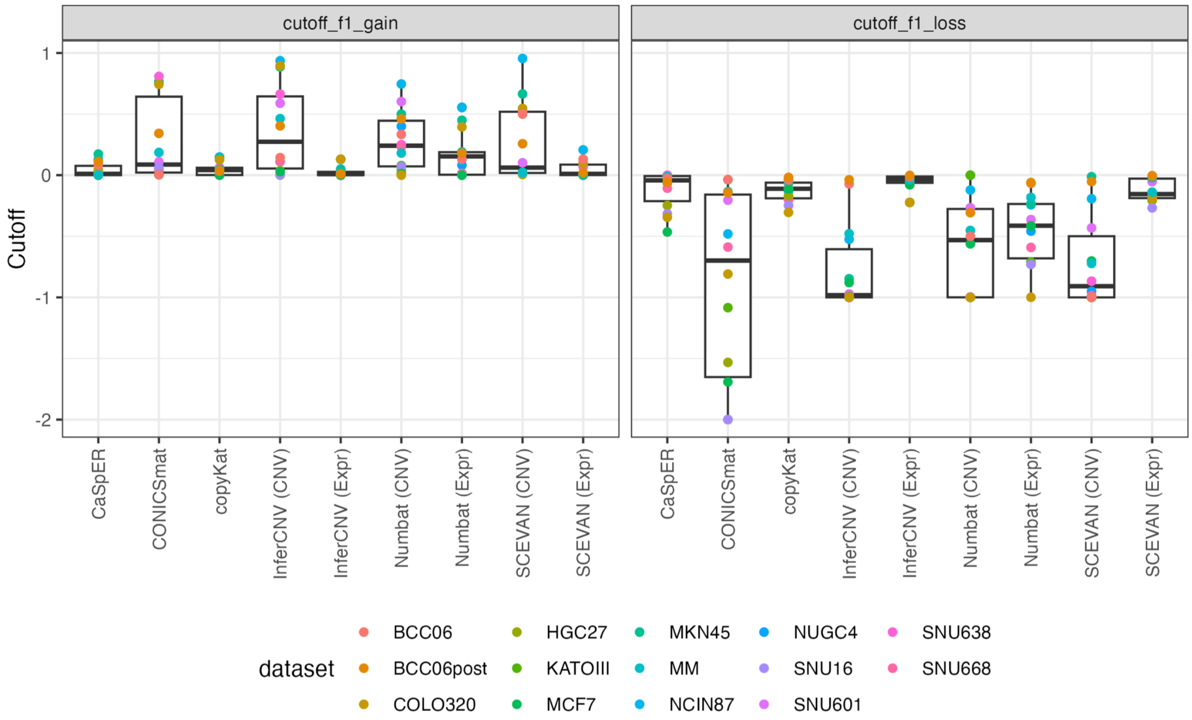


**Supplementary Figure 1.** Optimal gain and loss thresholds according to the multi-class F1 evaluation, after setting the baseline value of each method to 0.

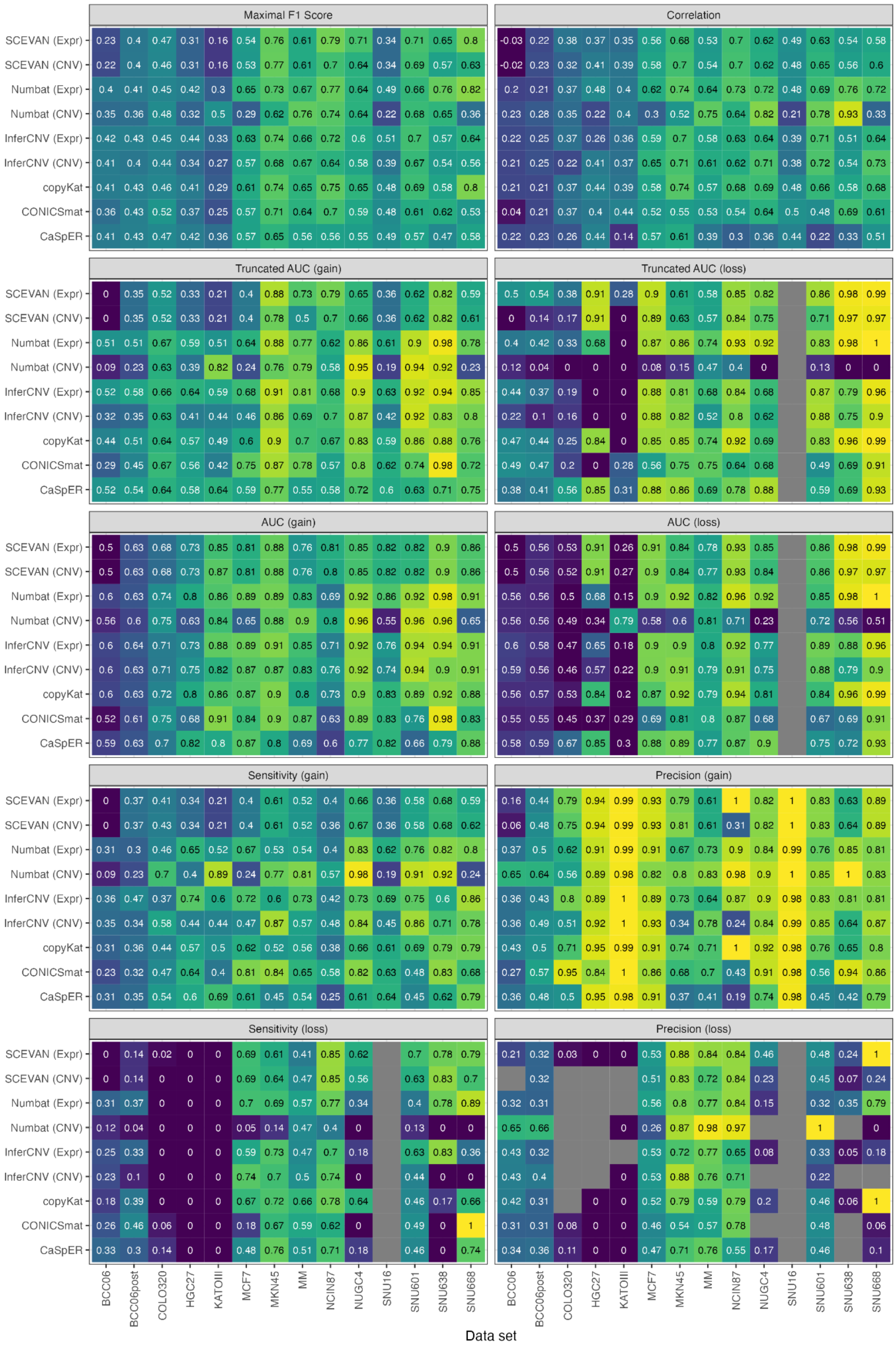


**Supplementary Figure 2.** Additional evaluation metrics for all datasets. The AUC values were calculated separately for gains vs all and loss vs all, same for the sensitivity and precision values. Here, the cutoffs for gains and losses were chosen based on maximal F1 scores.

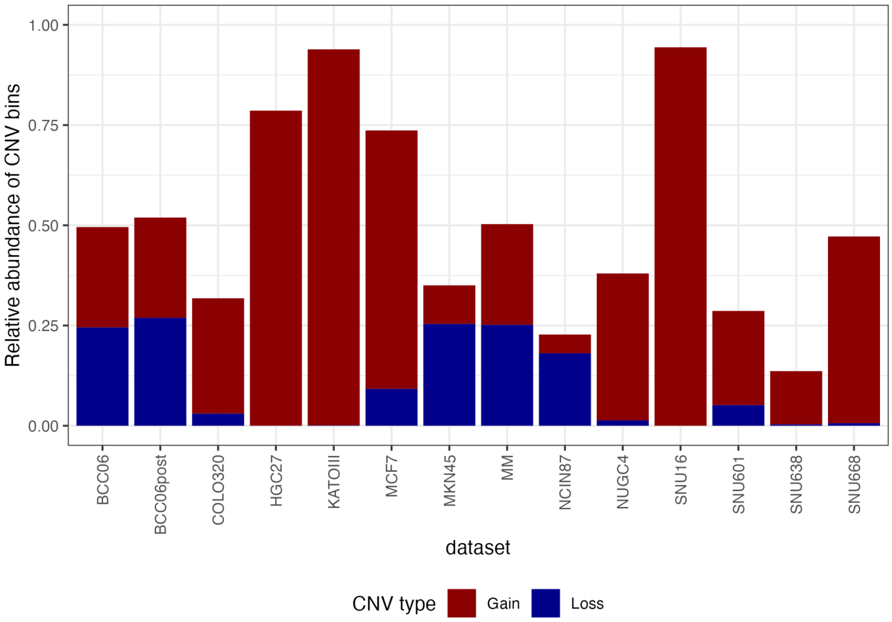


**Supplementary Figure 3.** CNV distribution across datasets

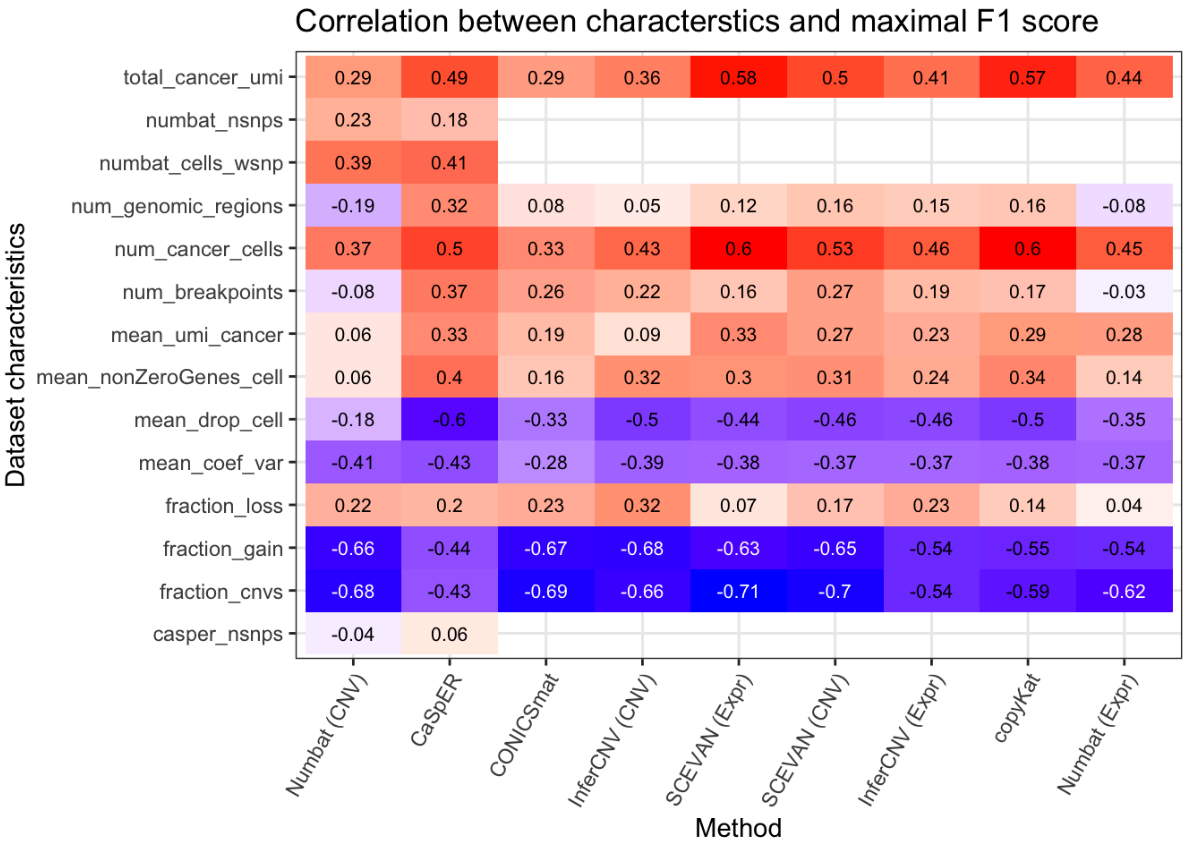


| | Numbat (CNV) | CaSpER | CONICSmat | InferCNV (CNV) | SCEVAN (Expr) | SCEVAN (CNV) | InferCNV (Expr) | copyKat | Numbat (Expr) |
|---|---|---|---|---|---|---|---|---|---|
| total_cancer_umi | 0.29 | 0.49 | 0.29 | 0.36 | 0.58 | 0.5 | 0.41 | 0.57 | 0.44 |
| numbat_nsnps | 0.23 | 0.18 | | | | | | | |
| numbat_cells_wsnp | 0.39 | 0.41 | | | | | | | |
| num_genomic_regions | -0.19 | 0.32 | 0.08 | 0.05 | 0.12 | 0.16 | 0.15 | 0.16 | -0.08 |
| num_cancer_cells | 0.37 | 0.5 | 0.33 | 0.43 | 0.6 | 0.53 | 0.46 | 0.6 | 0.45 |
| num_breakpoints | -0.08 | 0.37 | 0.26 | 0.22 | 0.16 | 0.27 | 0.19 | 0.17 | -0.03 |
| mean_umi_cancer | 0.06 | 0.33 | 0.19 | 0.09 | 0.33 | 0.27 | 0.23 | 0.29 | 0.28 |
| mean_nonZeroGenes_cell | 0.06 | 0.4 | 0.16 | 0.32 | 0.3 | 0.31 | 0.24 | 0.34 | 0.14 |
| mean_drop_cell | -0.18 | -0.6 | -0.33 | -0.5 | -0.44 | -0.46 | -0.46 | -0.5 | -0.35 |
| mean_coef_var | -0.41 | -0.43 | -0.28 | -0.39 | -0.38 | -0.37 | -0.37 | -0.38 | -0.37 |
| fraction_loss | 0.22 | 0.2 | 0.23 | 0.32 | 0.07 | 0.17 | 0.23 | 0.14 | 0.04 |
| fraction_gain | -0.66 | -0.44 | -0.67 | -0.68 | -0.63 | -0.65 | -0.54 | -0.55 | -0.54 |
| fraction_cnvs | -0.68 | -0.43 | -0.69 | -0.66 | -0.71 | -0.7 | -0.54 | -0.59 | -0.62 |
| casper_nsnps | -0.04 | 0.06 | | | | | | | |

**Supplementary Figure 4.** Correlation of different dataset characteristics with the prediction performance (maximal F1 score). Detailed description of all dataset characteristics found in Methods section.

**Supplementary Figure 5.** Karyograms of all tested datasets. See separate file (SupplementaryFigure5_karyograms.pdf)

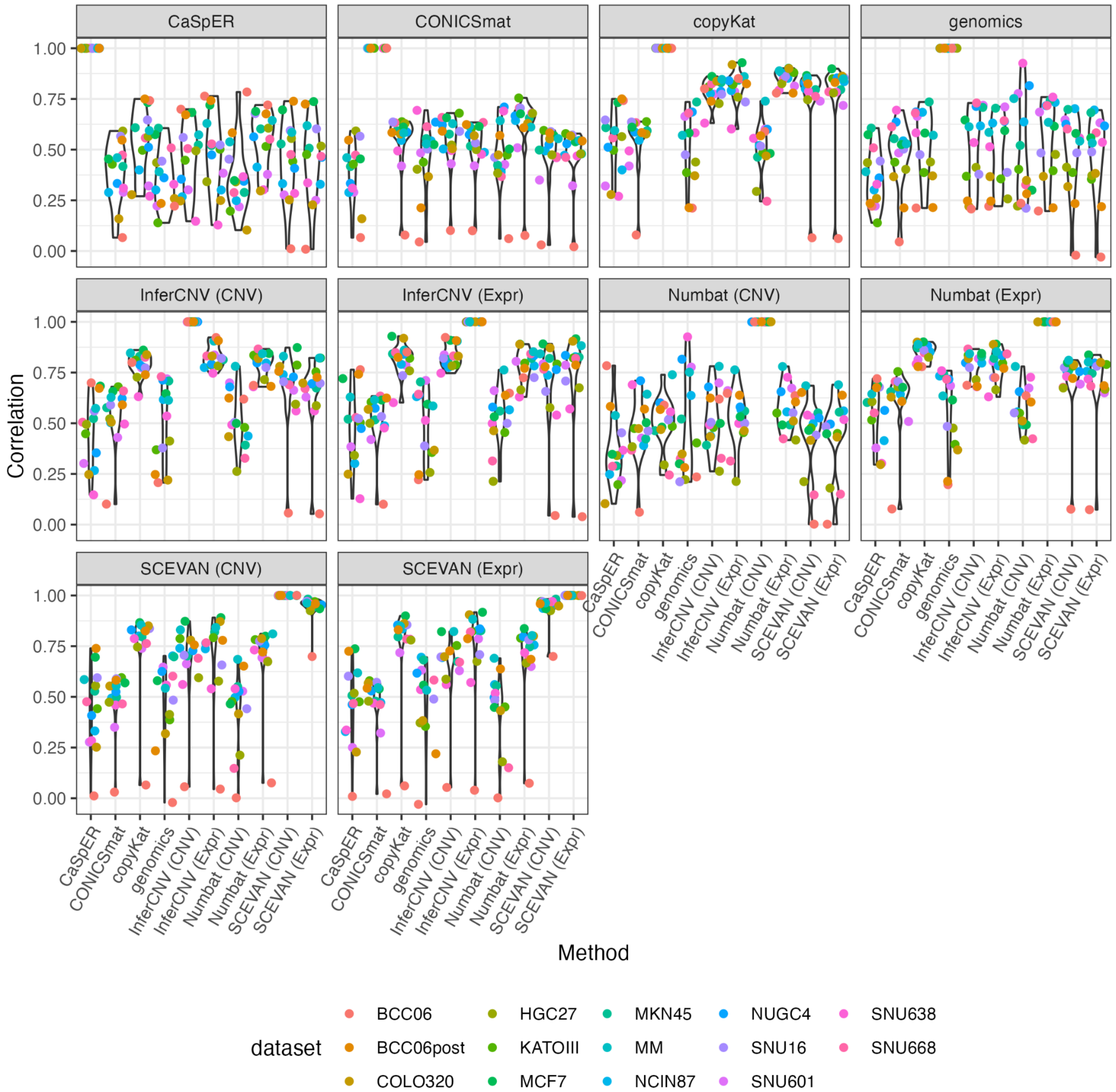


**Supplementary Figure 6.** Comparison of CNV predictions across scRNA-seq callers, quantified using Pearson correlation

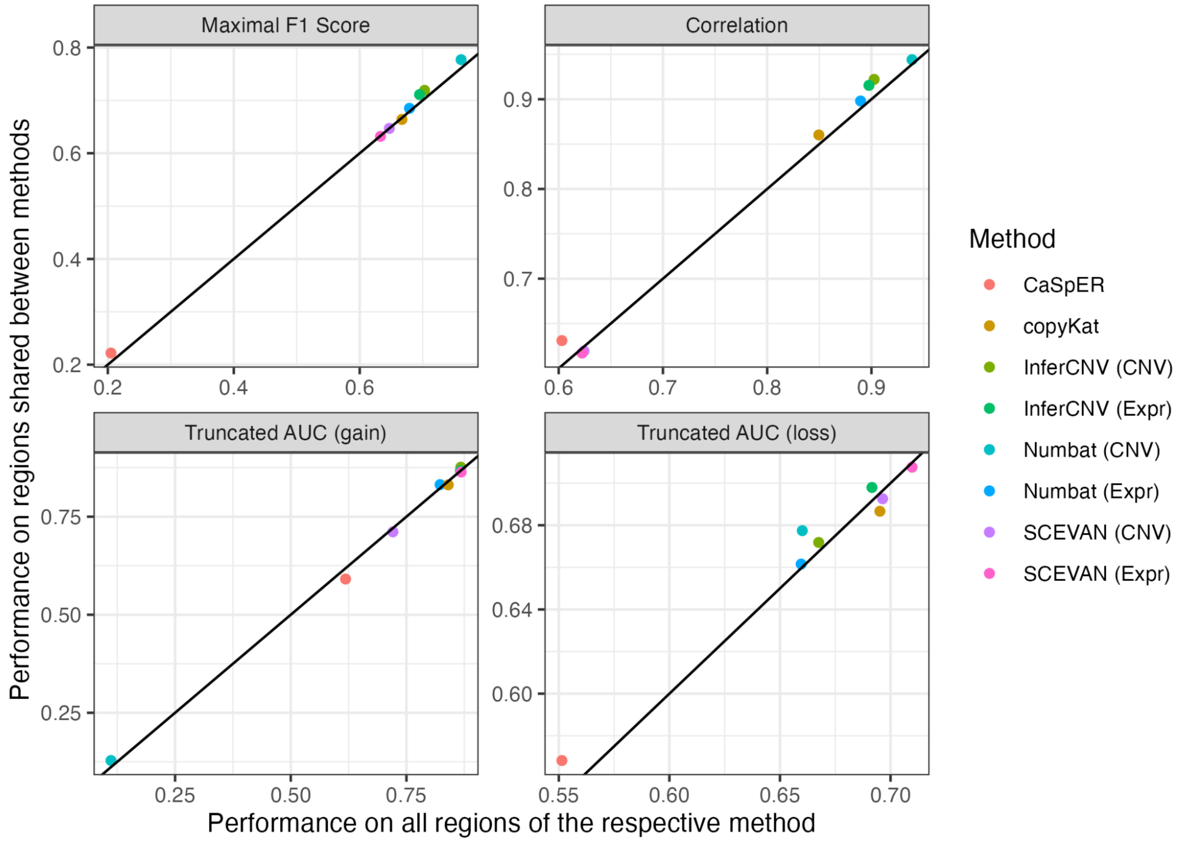


**Supplementary Figure 7.** Performance differences when looking into regions that are annotated by all methods (y axis) vs. all regions that are annotated by the respective method (x axis).

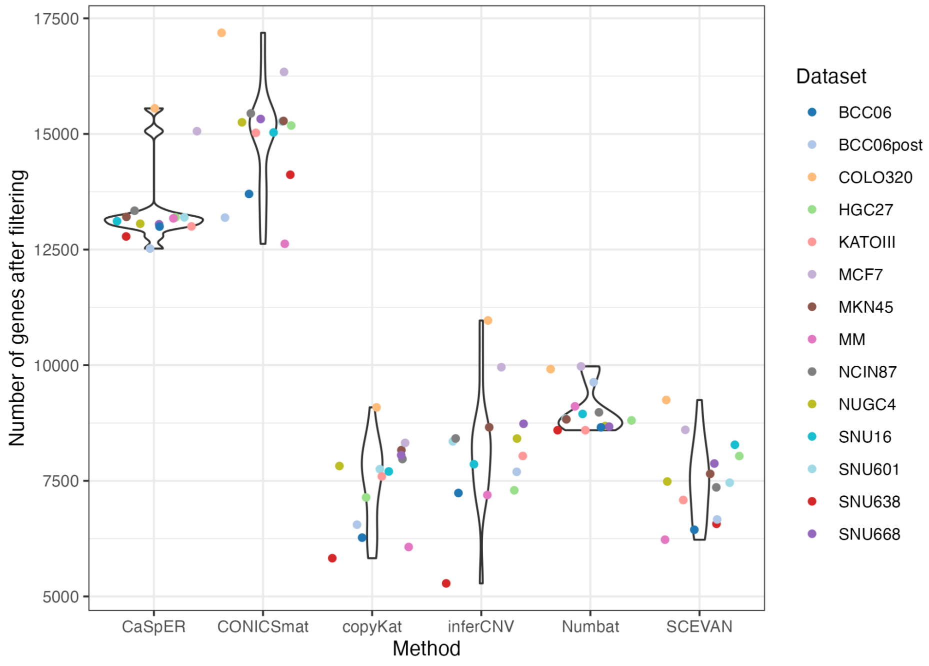


**Supplementary Figure 8.** Number of genes retained after filtering, for each method per dataset.

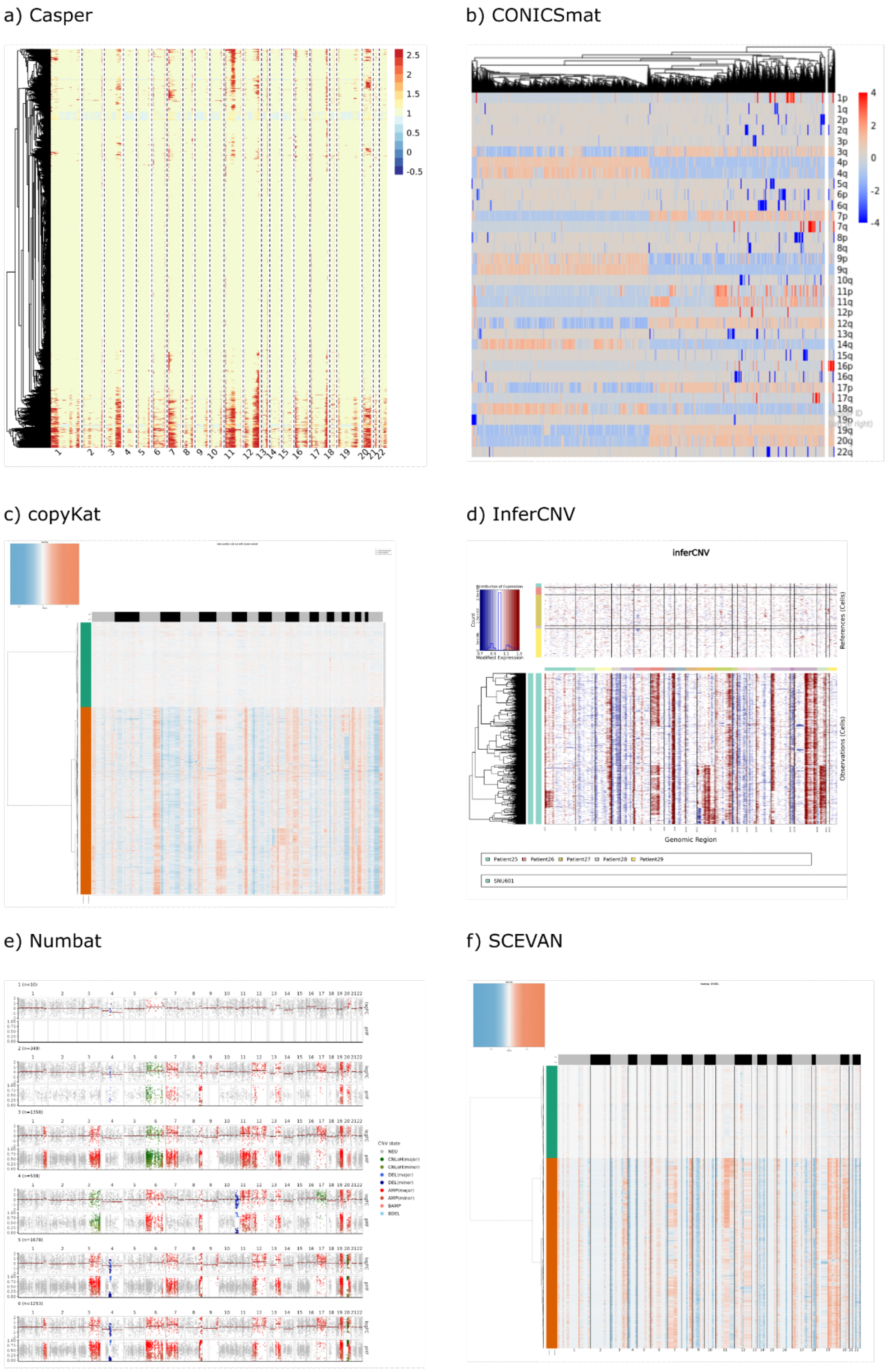


**Supplementary Figure 9.** Example result plots from each method for the SNU601 dataset (one representative plot was chosen per method).

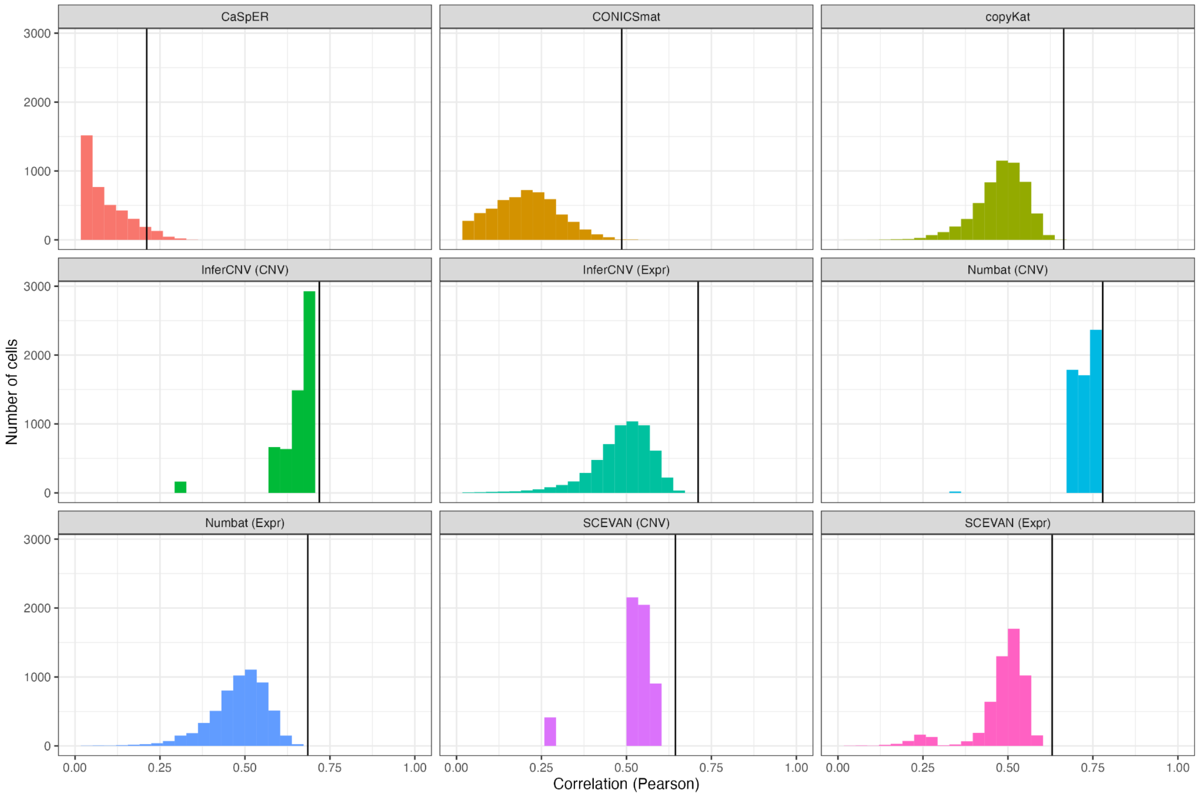


**Supplementary Figure 10.** Per-cell prediction results for all methods compared to the scWGS ground-truth for the SNU601 cell line. Vertical lines show the performance of the pseudobulk aggregated prediction. Correlation shown only for the value range 0-1 for better visibility, removing negative outliers.

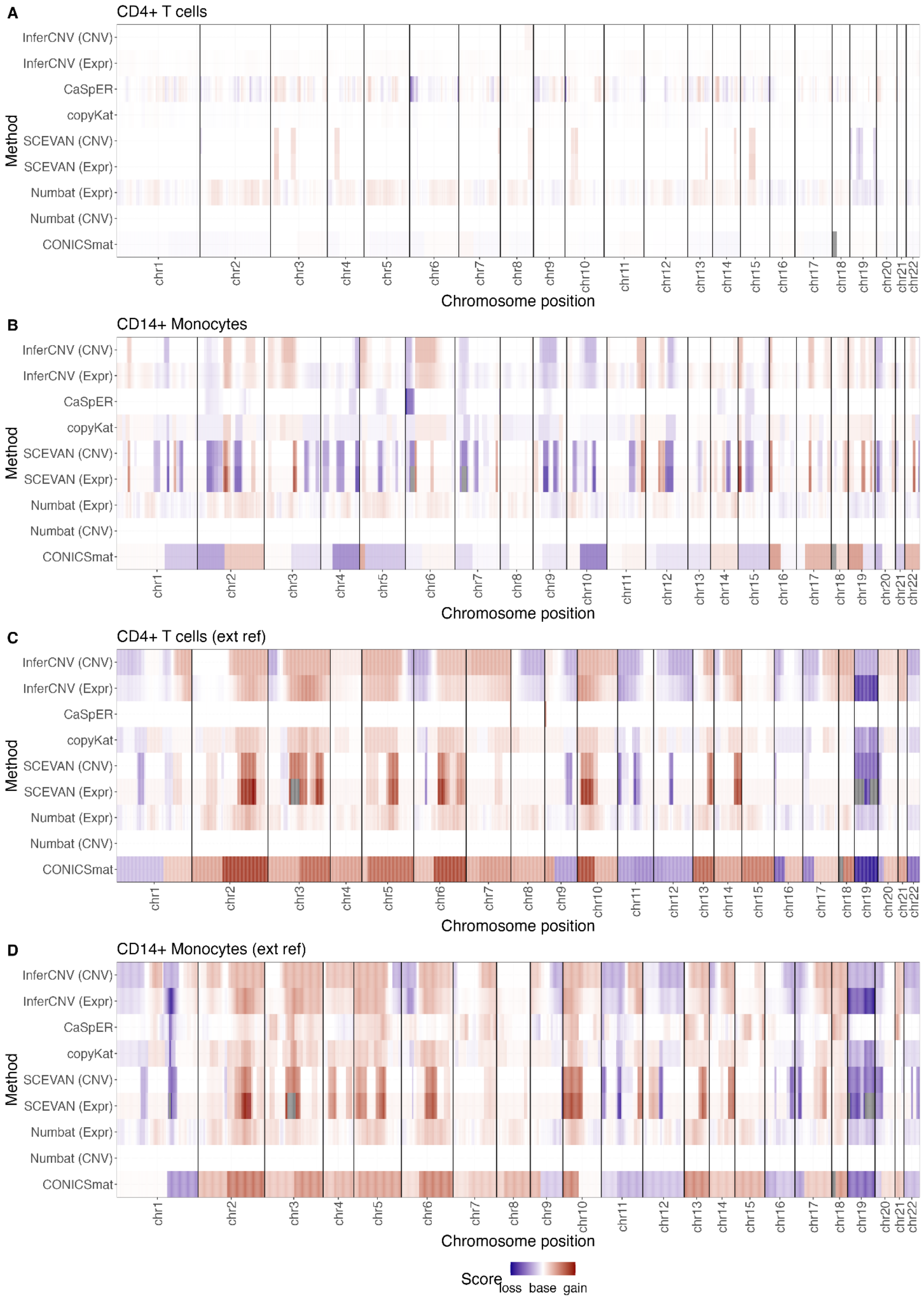


**Supplementary Figure 11.** Karyograms of CNVs in CD4+ T cells when using either CD4+ T cells (A) and CD14+ Monocytes (B) from the same dataset as reference cells, or CD4+ T cells (C) and CD14+ Monocytes (D) from an external dataset as reference.

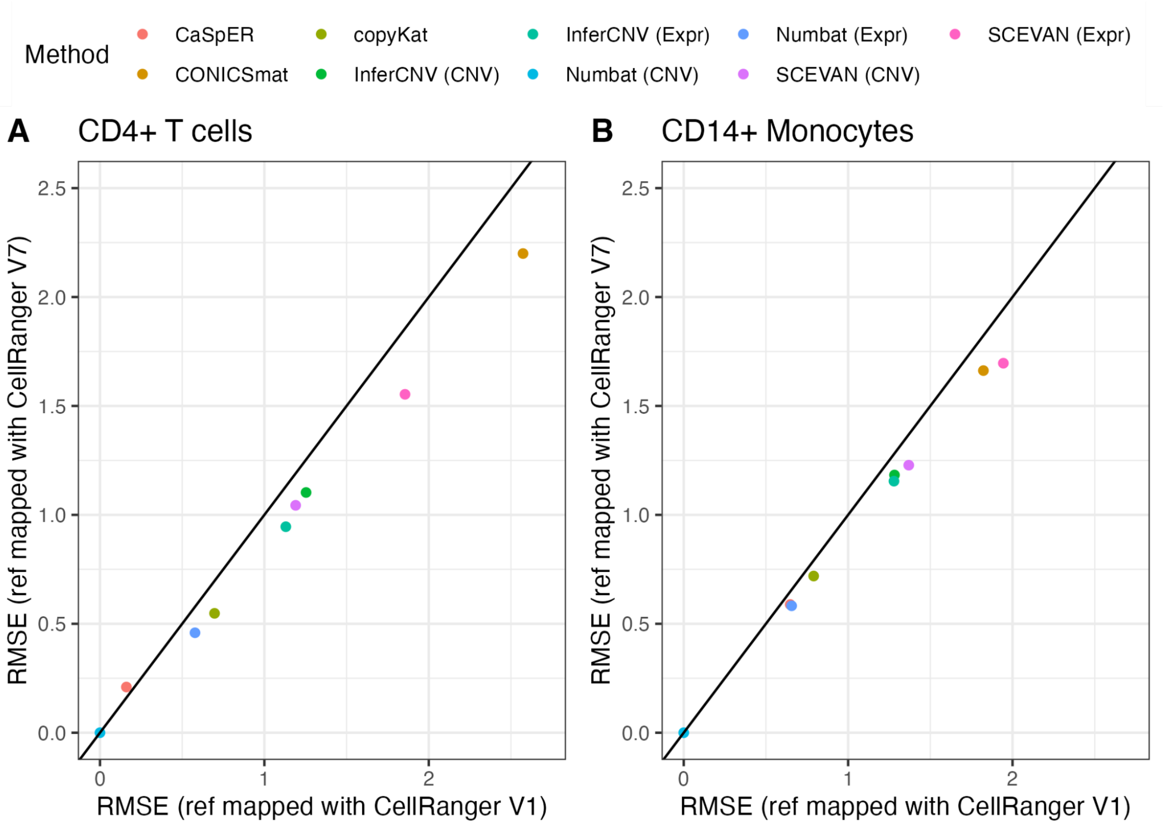


**Supplementary Figure 12.** Comparison of root mean square errors (RMSE) for identifying CNVs in a diploid dataset of CD4+ T cells when using two different mapper versions for the reference dataset: the reference contained CD4+ T cells (A) and CD14+ Monocytes (B), respectively, from a different study, once mapped with CellRanger version 1 and once with CellRanger version 7.

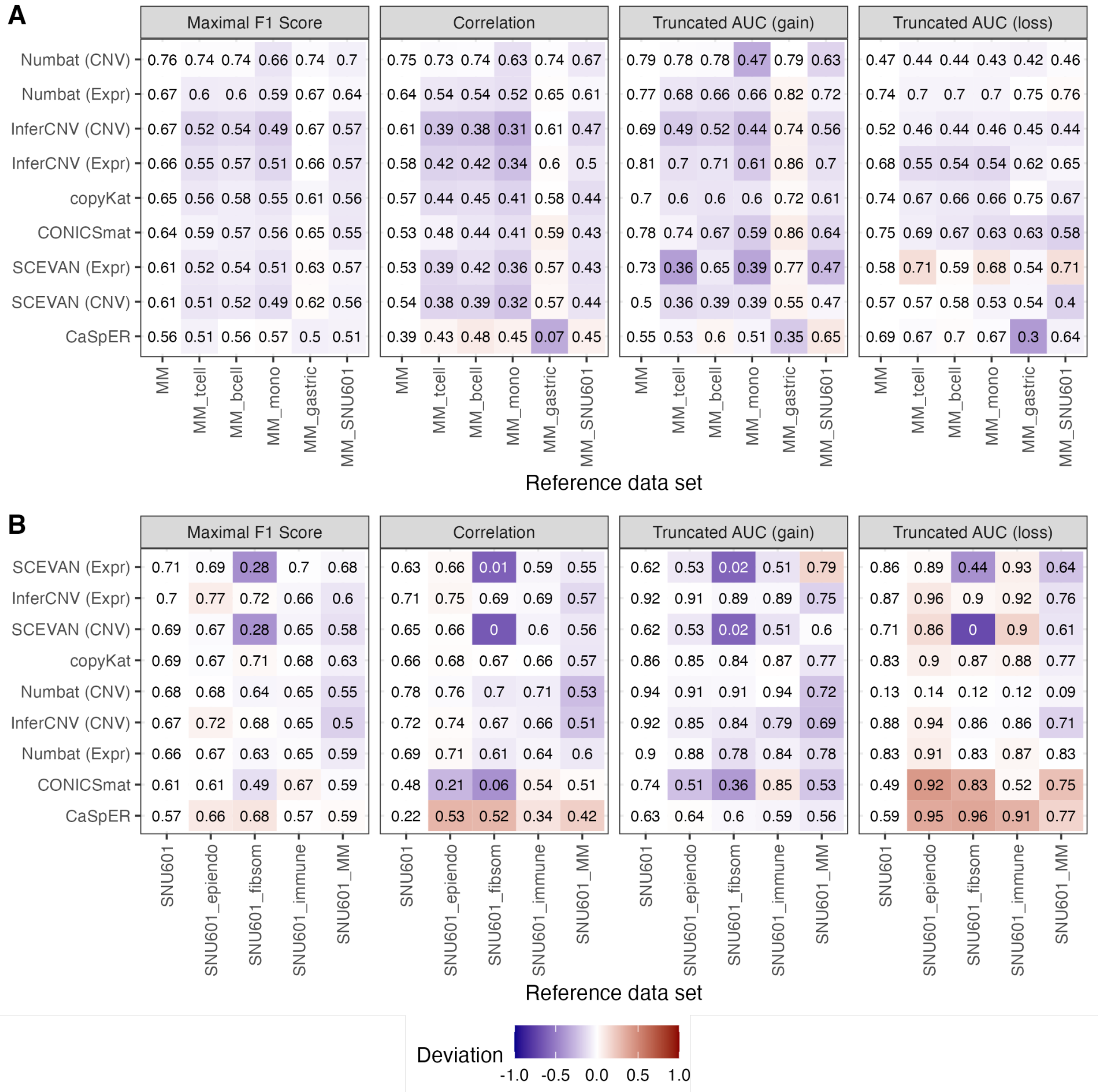


**Supplementary Figure 13.** Performance for predicting CNVs in the MM dataset (A) and the SNU601 dataset (B), when using different reference datasets.

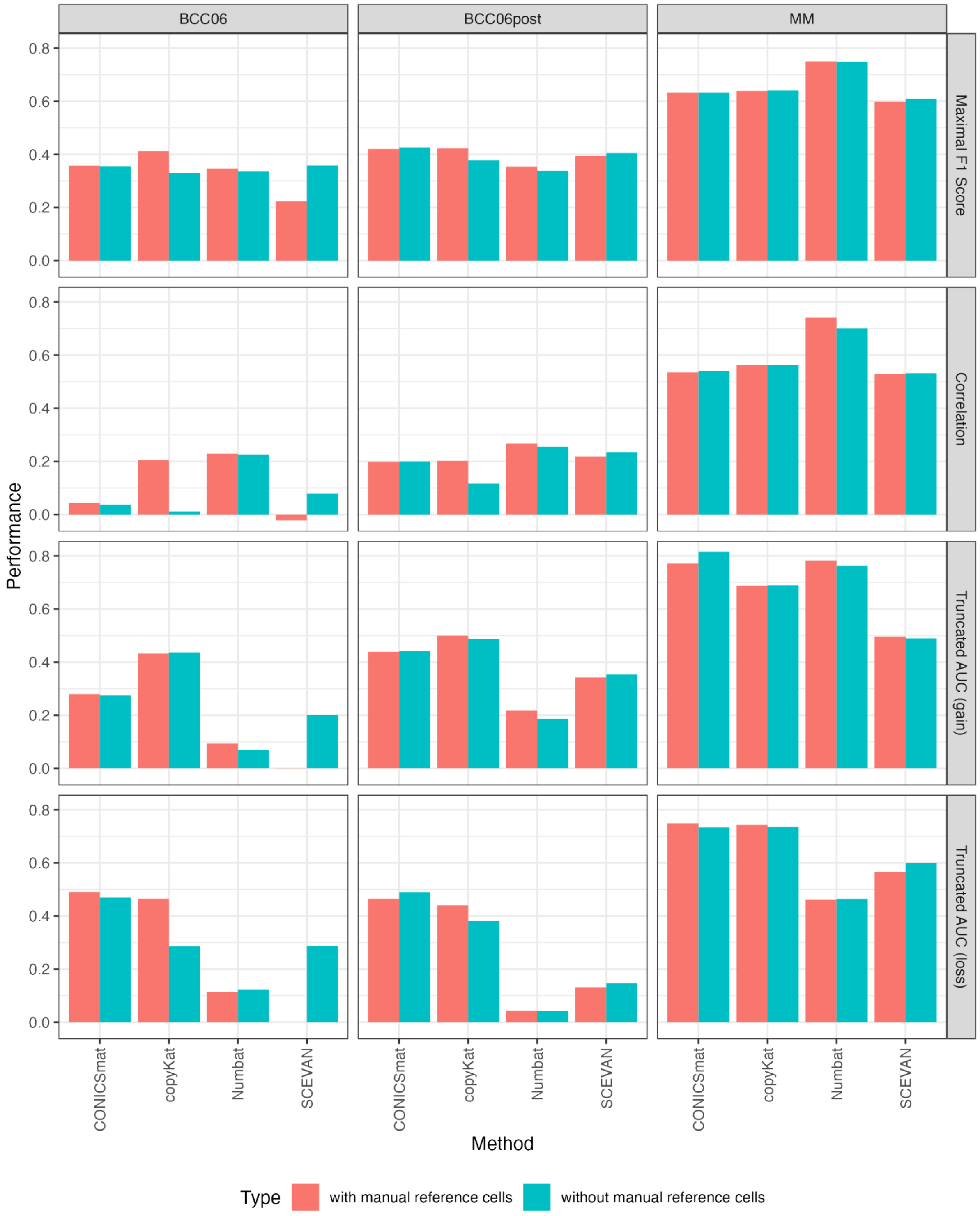


**Supplementary Figure 14.** Comparing CNV prediction performance when running copyKat, SCEVAN, Numbat and CONICSmat with and without manual reference cells. For the MM and the BCC06post samples, there is a high concordance of cell type annotations between the scenarios "with manual reference cells" and "without manual reference cells", which is very likely the explanation of why the performance between the two is not differing much.

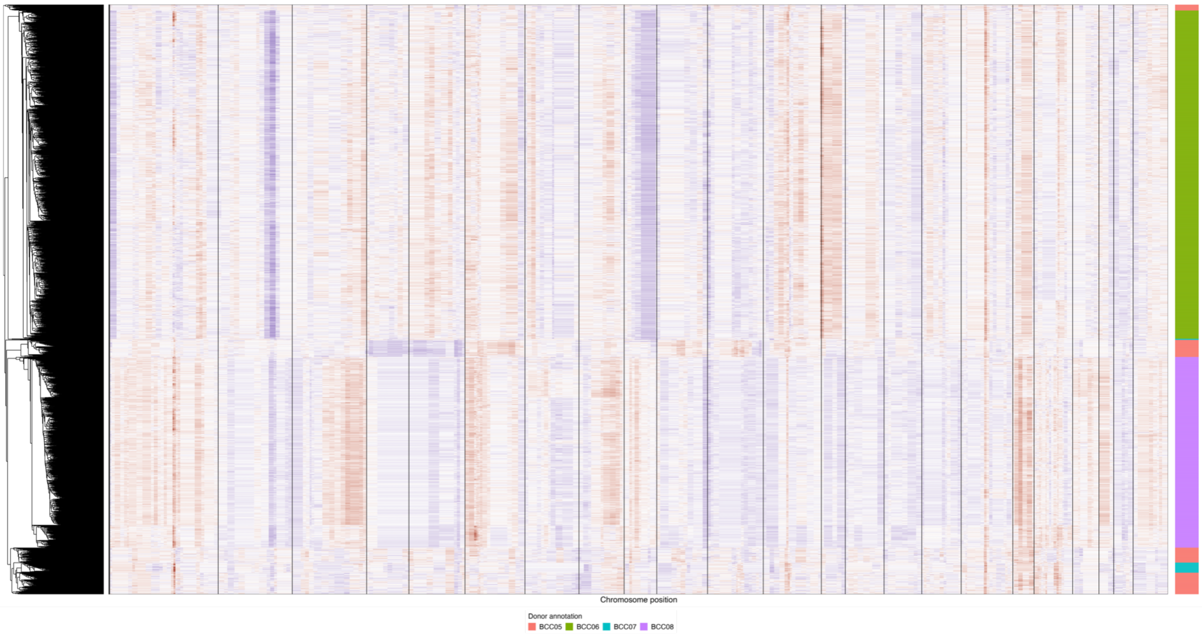


**Supplementary Figure 15.** Per cell CNV profile of the four BCC donors, estimated separately with copyKat and then clustered together afterwards.

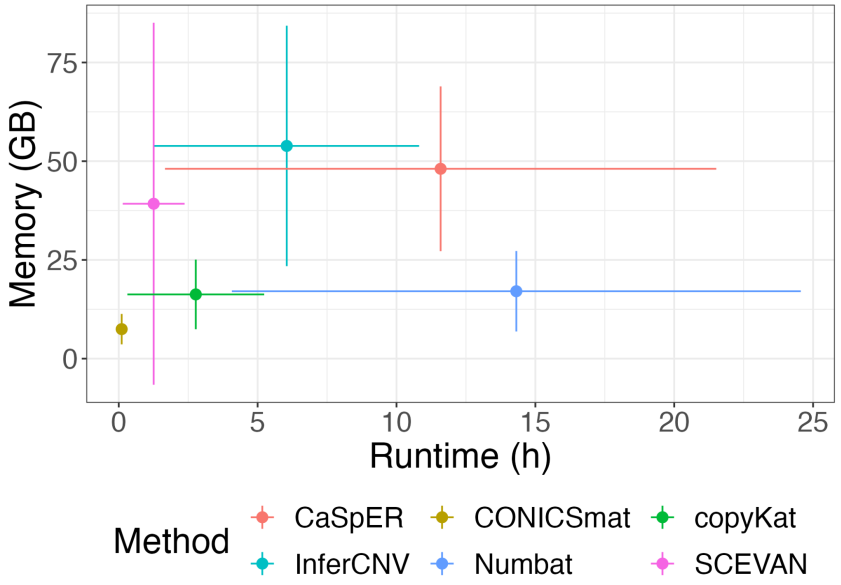


**Supplementary Figure 16.** Runtime and memory consumption of each method across all datasets, visualizing the mean with the dot size and minimum and maximum values with the horizontal and vertical lines.
