## Supplementary Figure 5 for "Benchmarking scRNA-seq copy number variation callers"

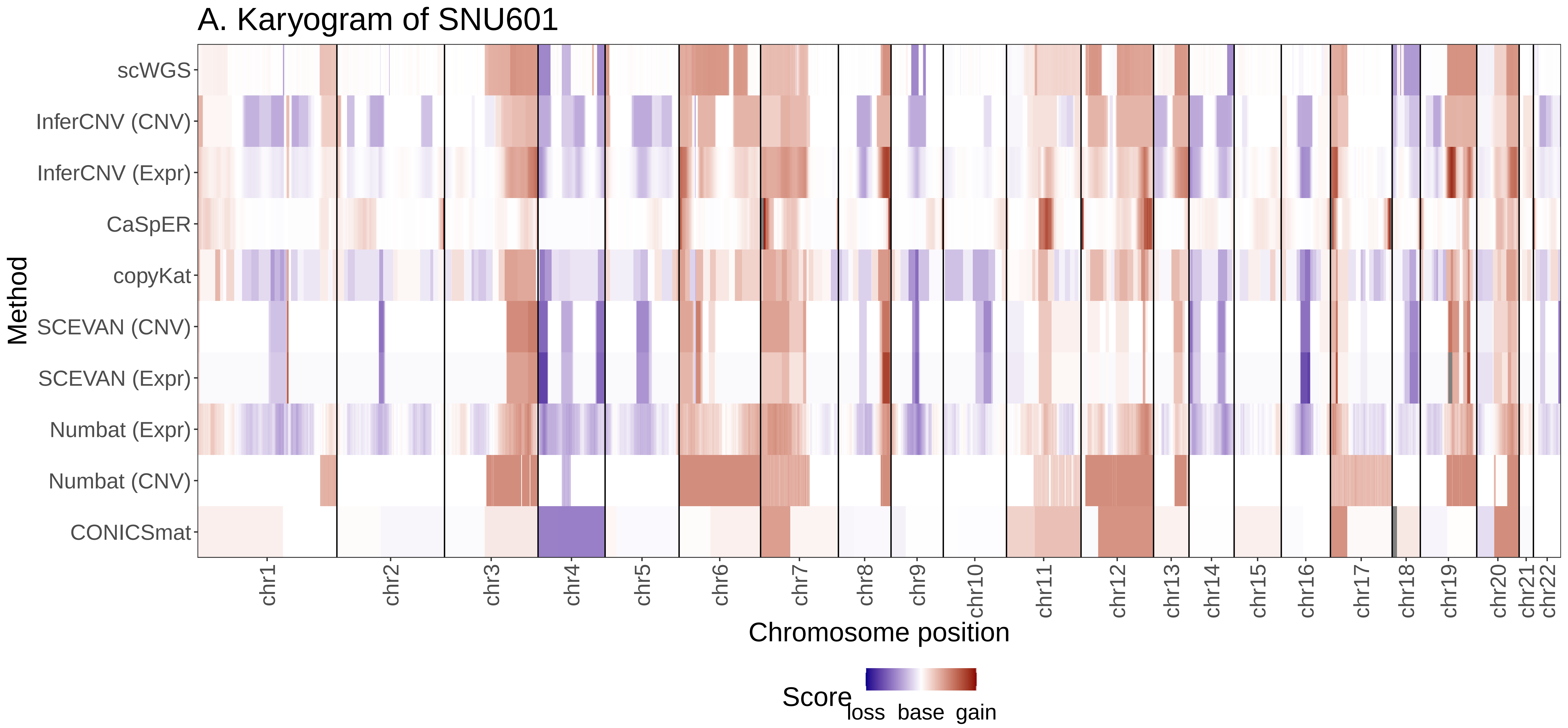

A. Karyogram of SNU601
Method
scWGS
InferCNV (CNV)
InferCNV (Expr)
CaSpER
copyKat
SCEVAN (CNV)
SCEVAN (Expr)
Numbat (Expr)
Numbat (CNV)
CONICSmat
chr1
chr2
chr3
chr4
chr5
chr6
chr7
chr8
chr9
chr10
chr11
chr12
chr13
chr14
chr15
chr16
chr17
chr18
chr19
chr20
chr21
chr22
Chromosome position
Score
loss base gain

B. Karyogram of NCIN87

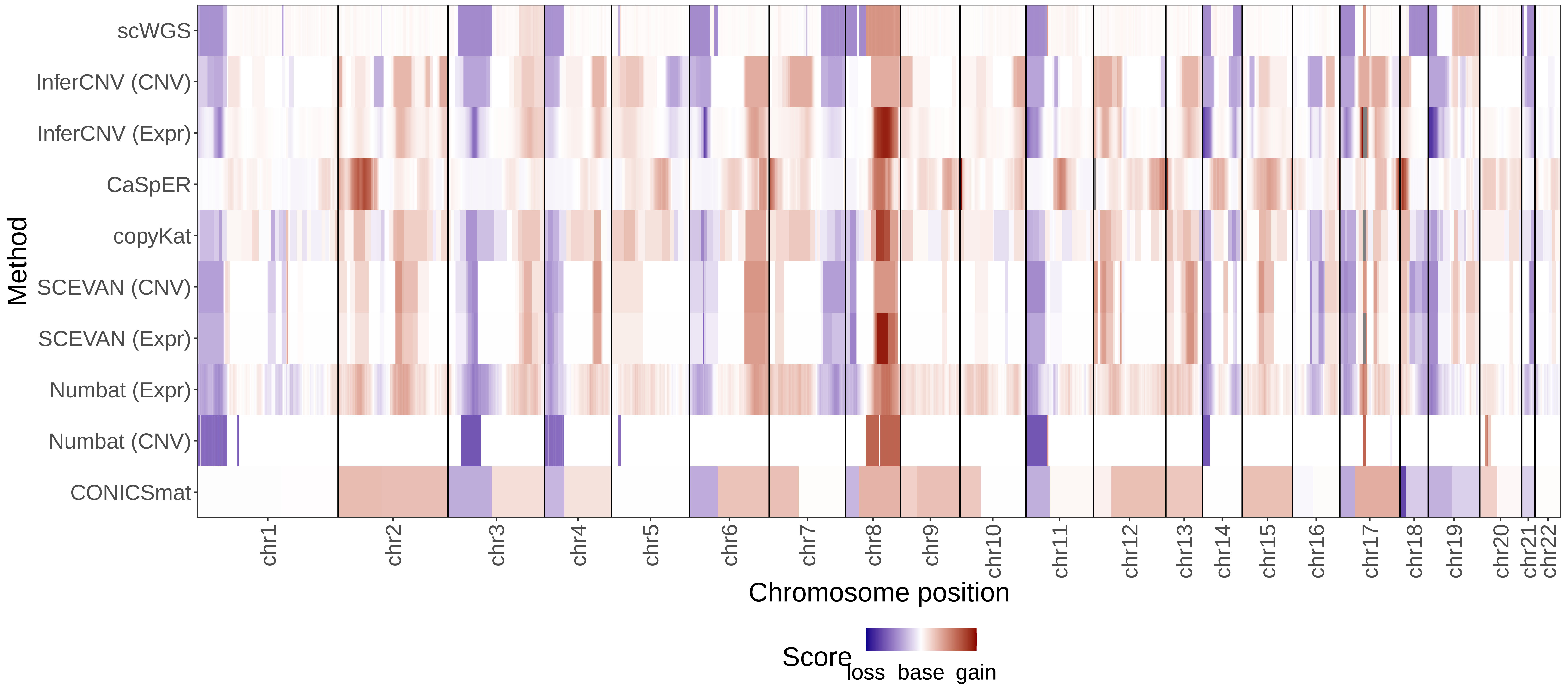

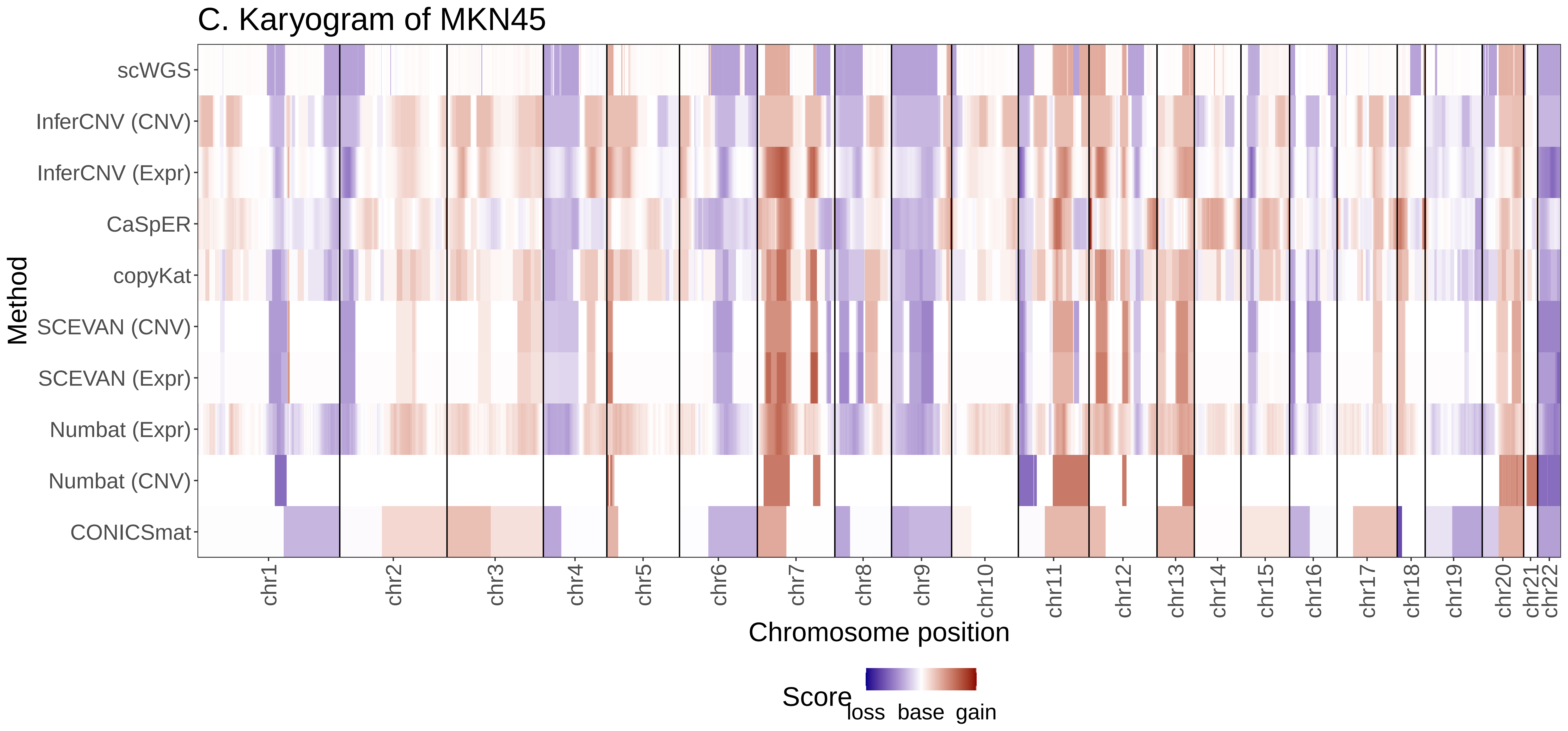

C. Karyogram of MKN45
scWGS
InferCNV (CNV)
InferCNV (Expr)
CaSpER
copyKat
SCEVAN (CNV)
SCEVAN (Expr)
Numbat (Expr)
Numbat (CNV)
CONICSmat
Method
chr1
chr2
chr3
chr4
chr5
chr6
chr7
chr8
chr9
chr10
chr11
chr12
chr13
chr14
chr15
chr16
chr17
chr18
chr19
chr20
chr21
chr22
Chromosome position
Score
loss base gain

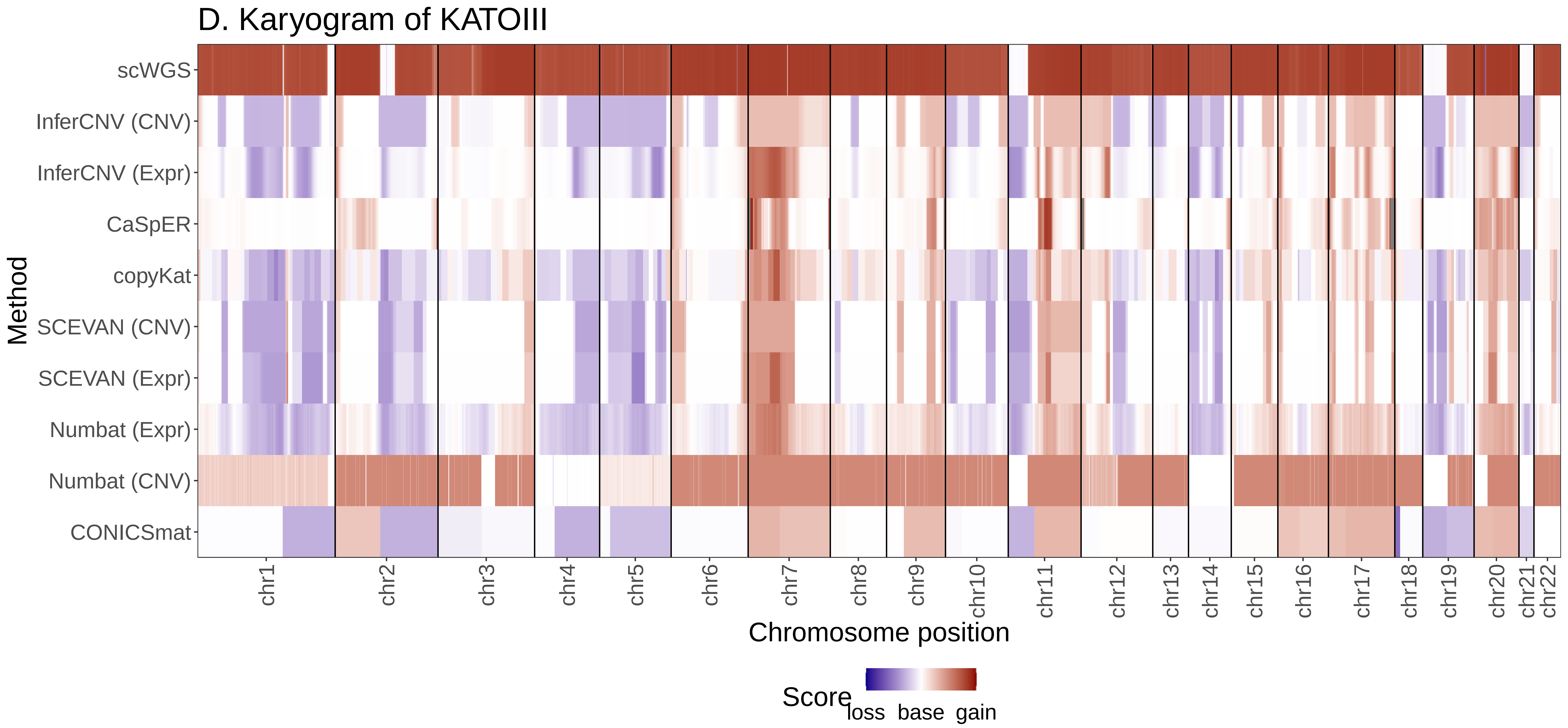
D. Karyogram of KATOIII
Method
scWGS
InferCNV (CNV)
InferCNV (Expr)
CaSpER
copyKat
SCEVAN (CNV)
SCEVAN (Expr)
Numbat (Expr)
Numbat (CNV)
CONICSmat
chr1
chr2
chr3
chr4
chr5
chr6
chr7
chr8
chr9
chr10
chr11
chr12
chr13
chr14
chr15
chr16
chr17
chr18
chr19
chr20
chr21
chr22
Chromosome position
Score
loss base gain

### E. Karyogram of NUGC4

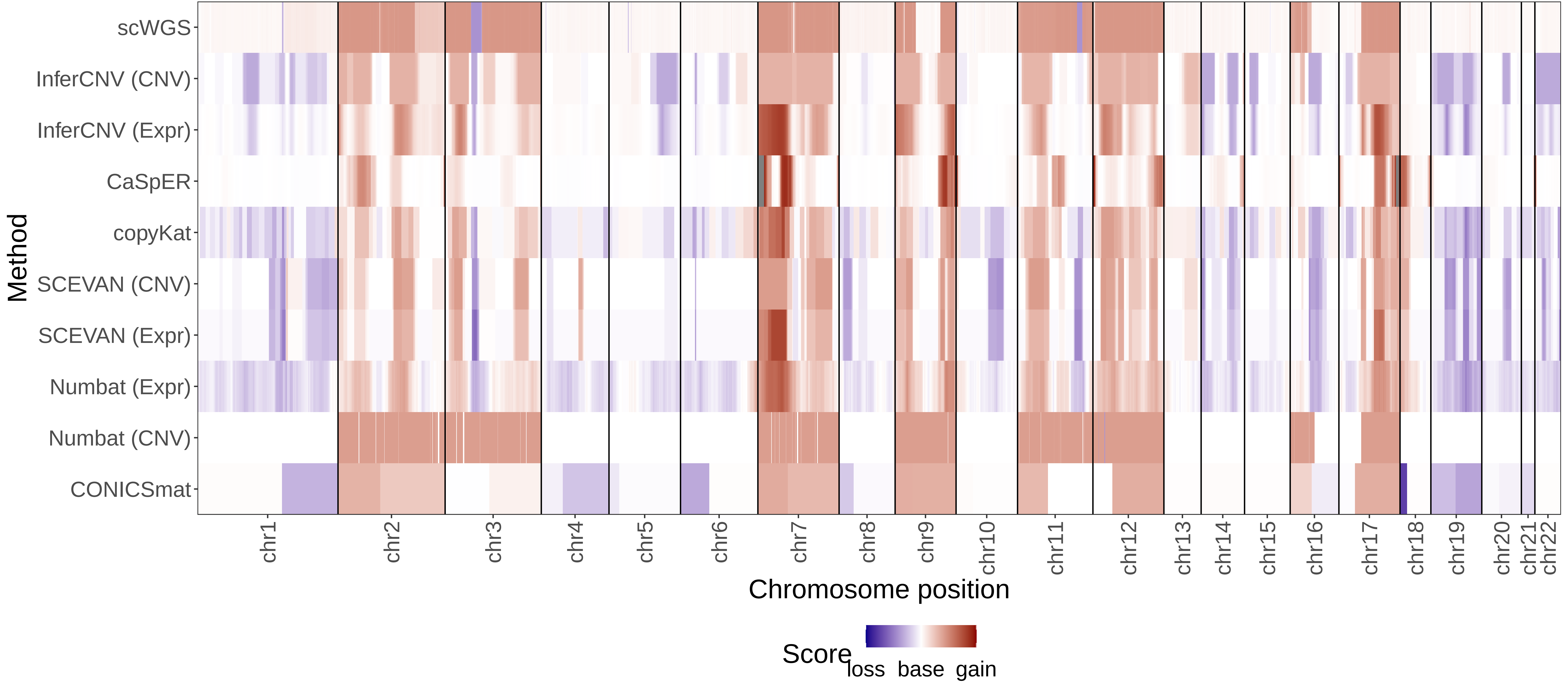

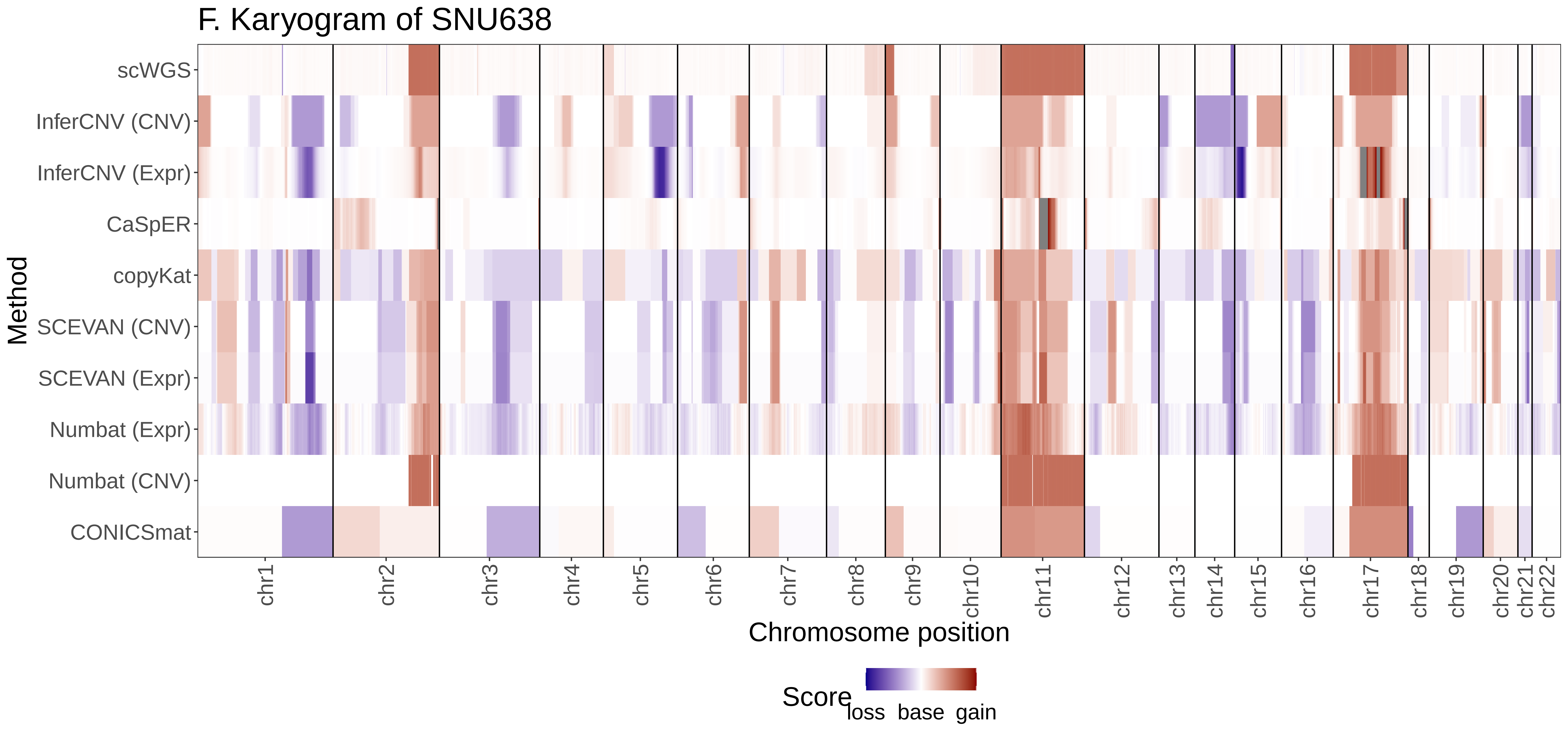
F. Karyogram of SNU638
Method
scWGS
InferCNV (CNV)
InferCNV (Expr)
CaSpER
copyKat
SCEVAN (CNV)
SCEVAN (Expr)
Numbat (Expr)
Numbat (CNV)
CONICSmat
chr1
chr2
chr3
chr4
chr5
chr6
chr7
chr8
chr9
chr10
chr11
chr12
chr13
chr14
chr15
chr16
chr17
chr18
chr19
chr20
chr21
chr22
Chromosome position
Score
loss
base
gain

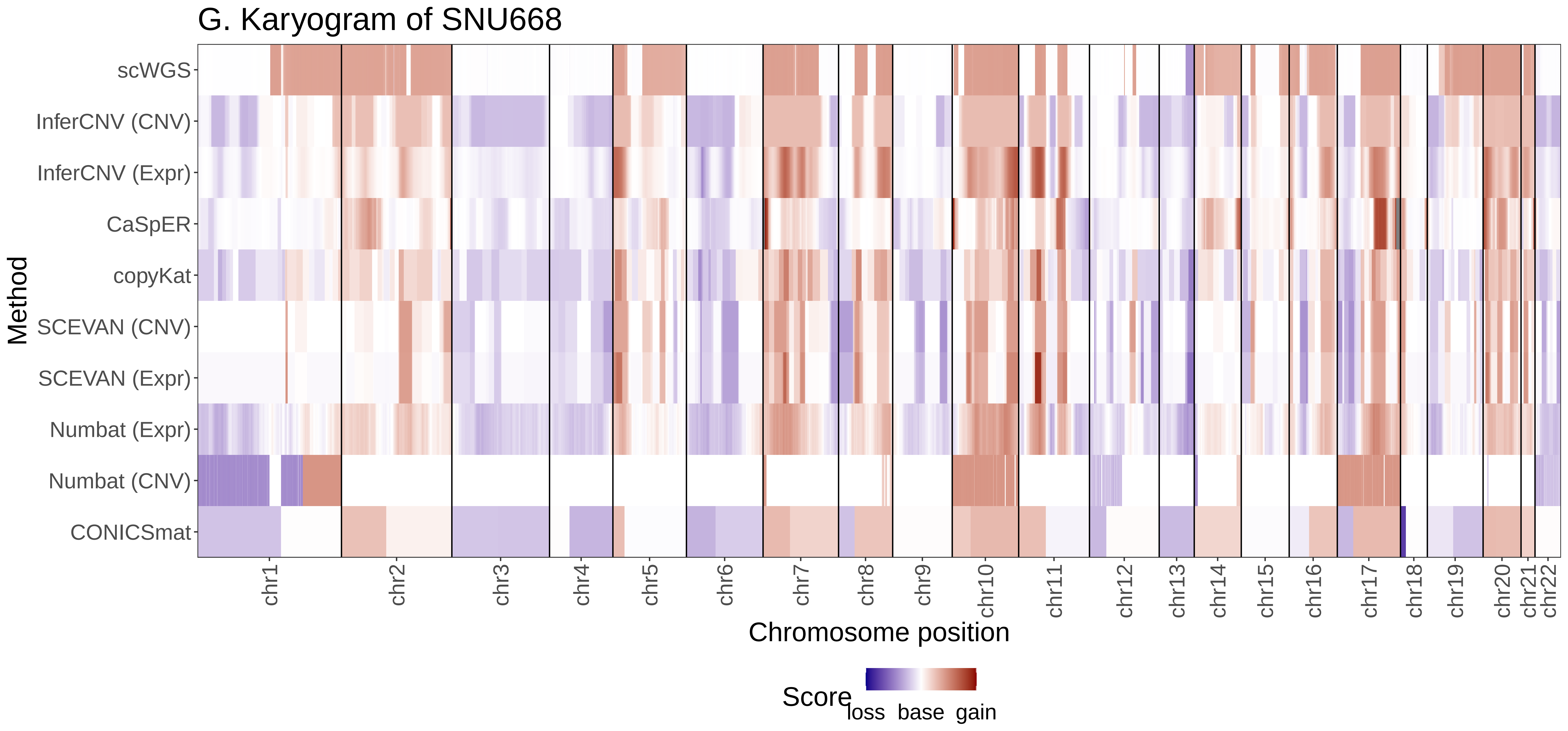
G. Karyogram of SNU668
Method
scWGS
InferCNV (CNV)
InferCNV (Expr)
CaSpER
copyKat
SCEVAN (CNV)
SCEVAN (Expr)
Numbat (Expr)
Numbat (CNV)
CONICSmat
chr1
chr2
chr3
chr4
chr5
chr6
chr7
chr8
chr9
chr10
chr11
chr12
chr13
chr14
chr15
chr16
chr17
chr18
chr19
chr20
chr21
chr22
Chromosome position
Score
loss base gain

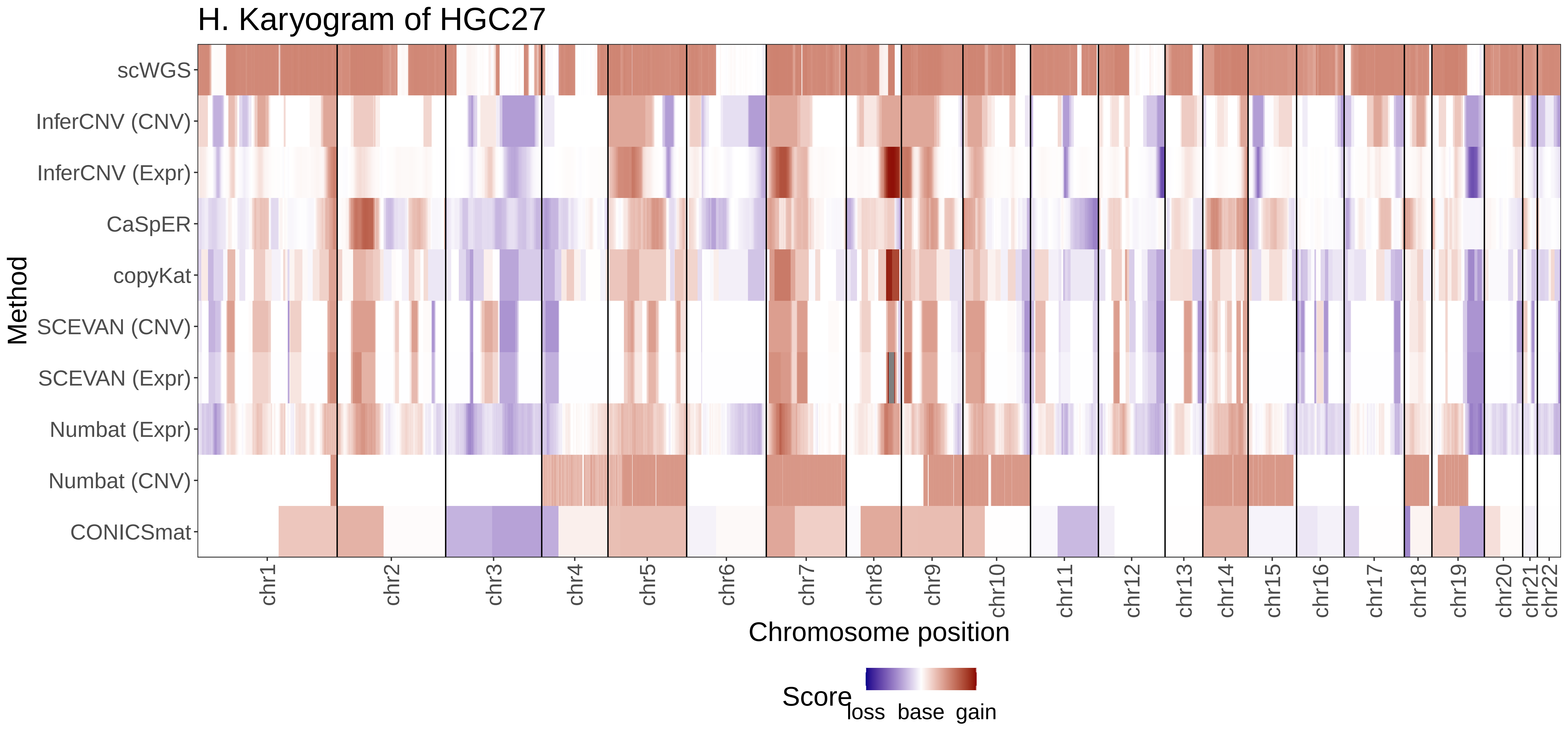
H. Karyogram of HGC27
scWGS
InferCNV (CNV)
InferCNV (Expr)
CaSpER
copyKat
SCEVAN (CNV)
SCEVAN (Expr)
Numbat (Expr)
Numbat (CNV)
CONICSmat
Method
chr1
chr2
chr3
chr4
chr5
chr6
chr7
chr8
chr9
chr10
chr11
chr12
chr13
chr14
chr15
chr16
chr17
chr18
chr19
chr20
chr21
chr22
Chromosome position
Score
loss
base
gain

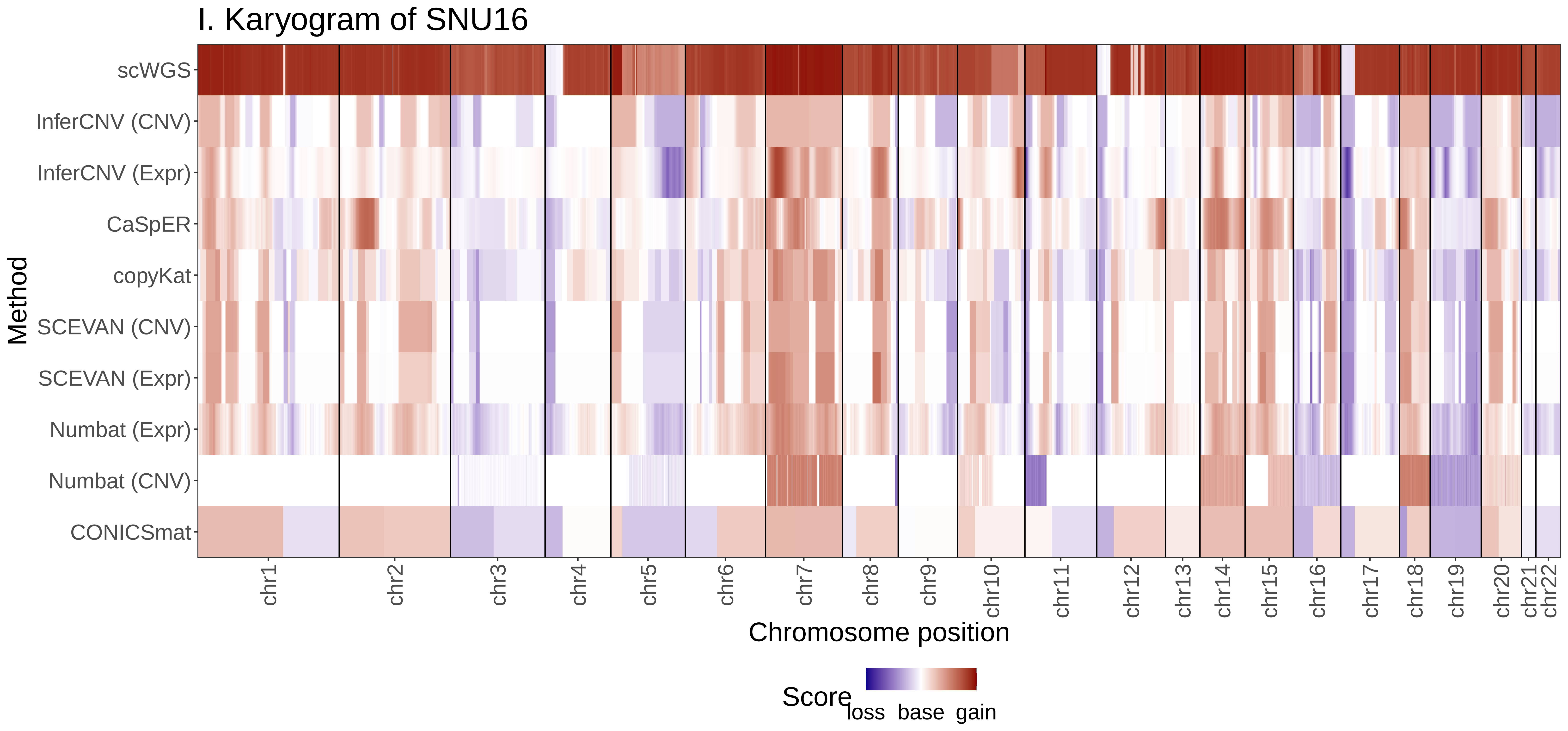

I. Karyogram of SNU16
Method
scWGS
InferCNV (CNV)
InferCNV (Expr)
CaSpER
copyKat
SCEVAN (CNV)
SCEVAN (Expr)
Numbat (Expr)
Numbat (CNV)
CONICSmat
chr1
chr2
chr3
chr4
chr5
chr6
chr7
chr8
chr9
chr10
chr11
chr12
chr13
chr14
chr15
chr16
chr17
chr18
chr19
chr20
chr21
chr22
Chromosome position
Score
loss
base
gain

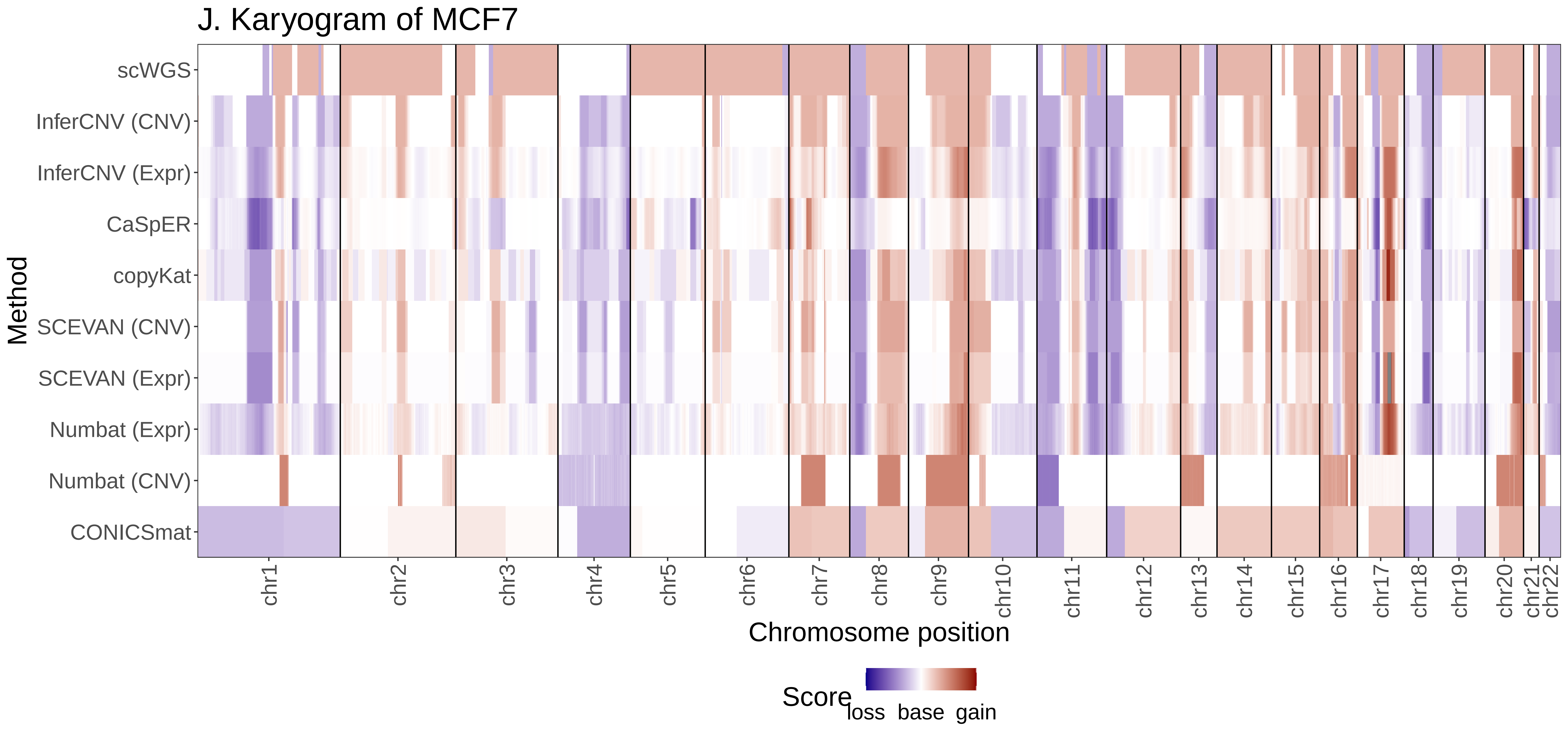

J. Karyogram of MCF7
Method
scWGS
InferCNV (CNV)
InferCNV (Expr)
CaSpER
copyKat
SCEVAN (CNV)
SCEVAN (Expr)
Numbat (Expr)
Numbat (CNV)
CONICSmat
chr1
chr2
chr3
chr4
chr5
chr6
chr7
chr8
chr9
chr10
chr11
chr12
chr13
chr14
chr15
chr16
chr17
chr18
chr19
chr20
chr21
chr22
Chromosome position
Score
loss
base
gain

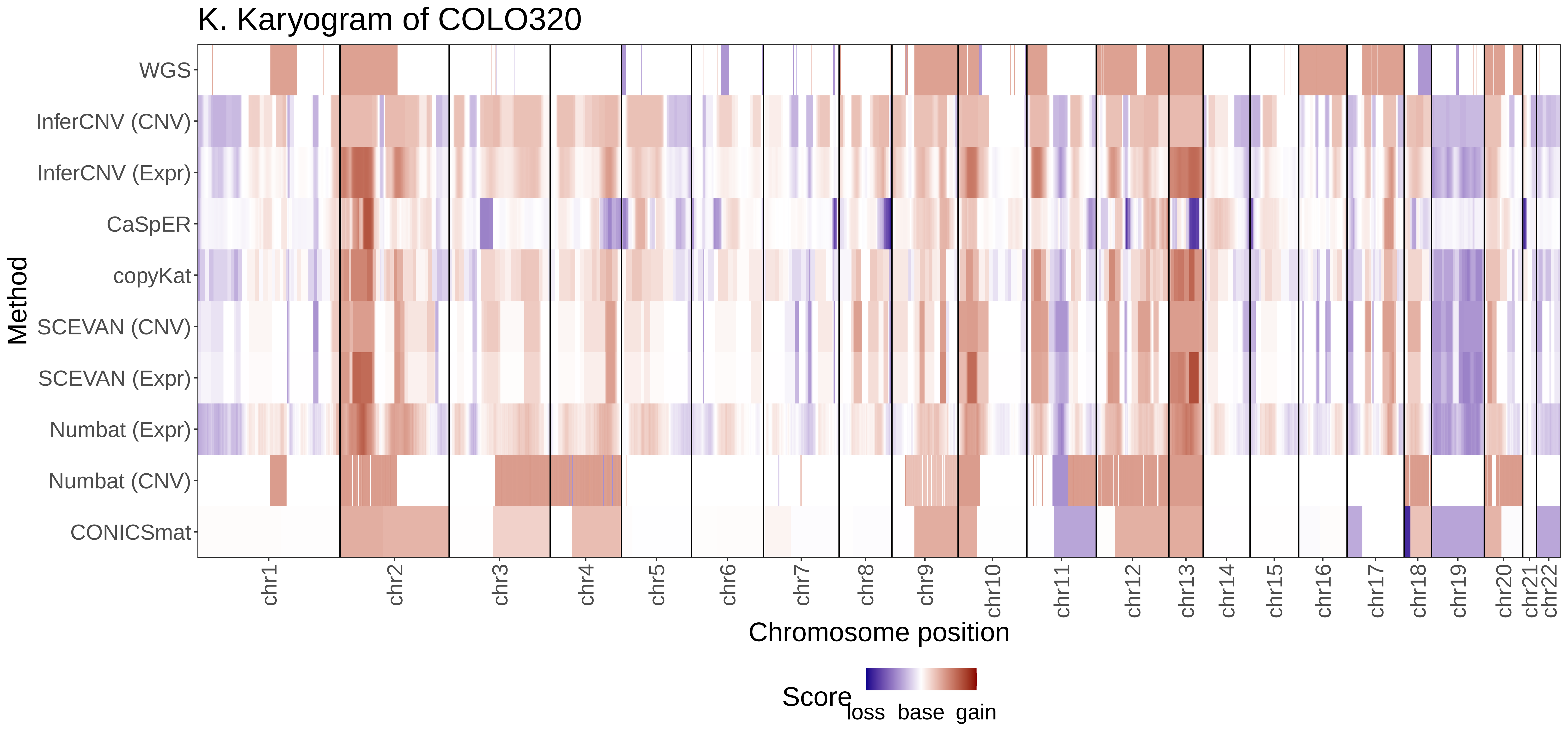

K. Karyogram of COLO320
Method
WGS
InferCNV (CNV)
InferCNV (Expr)
CaSpER
copyKat
SCEVAN (CNV)
SCEVAN (Expr)
Numbat (Expr)
Numbat (CNV)
CONICSmat
chr1
chr2
chr3
chr4
chr5
chr6
chr7
chr8
chr9
chr10
chr11
chr12
chr13
chr14
chr15
chr16
chr17
chr18
chr19
chr20
chr21
chr22
Chromosome position
Score
loss
base
gain

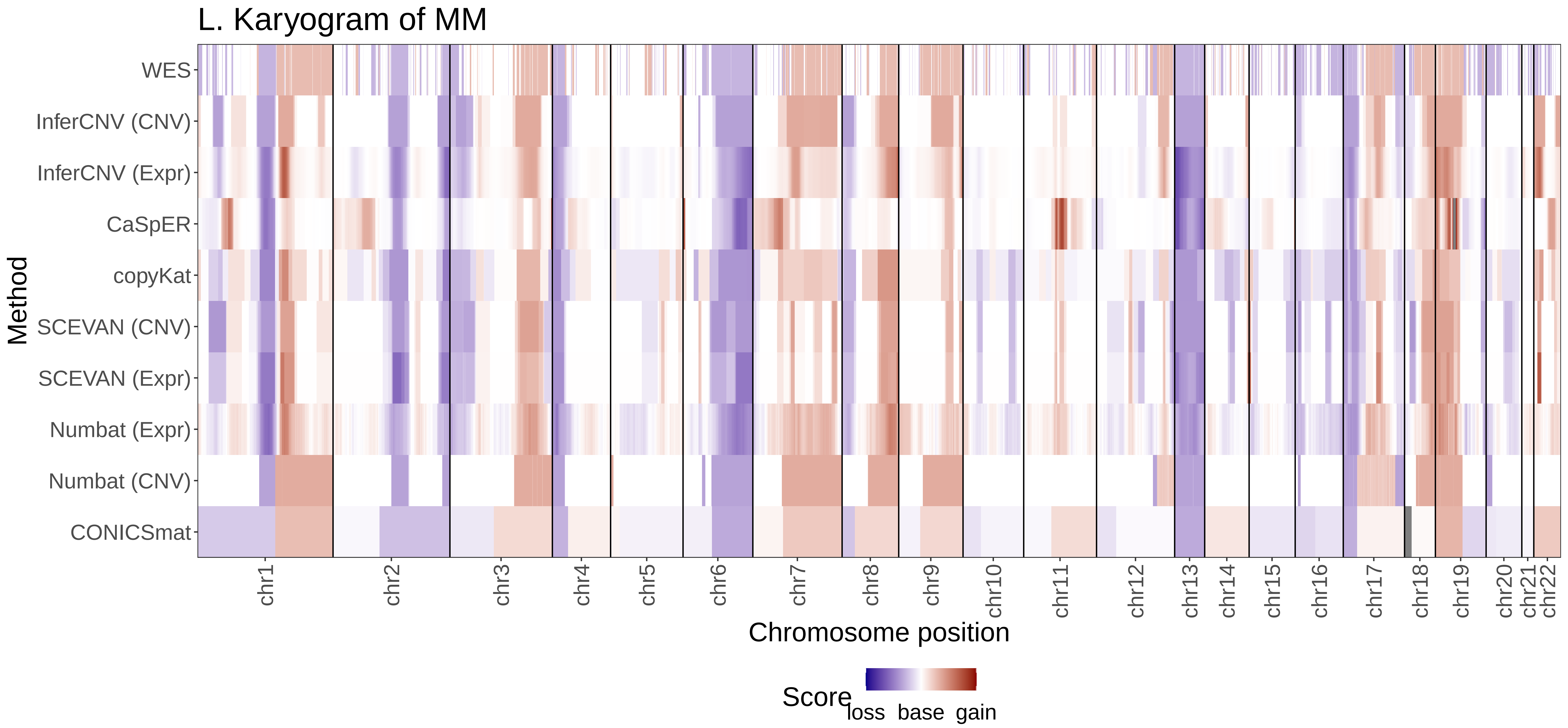

L. Karyogram of MM
WES
InferCNV (CNV)
InferCNV (Expr)
CaSpER
copyKat
SCEVAN (CNV)
SCEVAN (Expr)
Numbat (Expr)
Numbat (CNV)
CONICSmat
Method
chr1
chr2
chr3
chr4
chr5
chr6
chr7
chr8
chr9
chr10
chr11
chr12
chr13
chr14
chr15
chr16
chr17
chr18
chr19
chr20
chr21
chr22
Chromosome position
Score
loss
base
gain

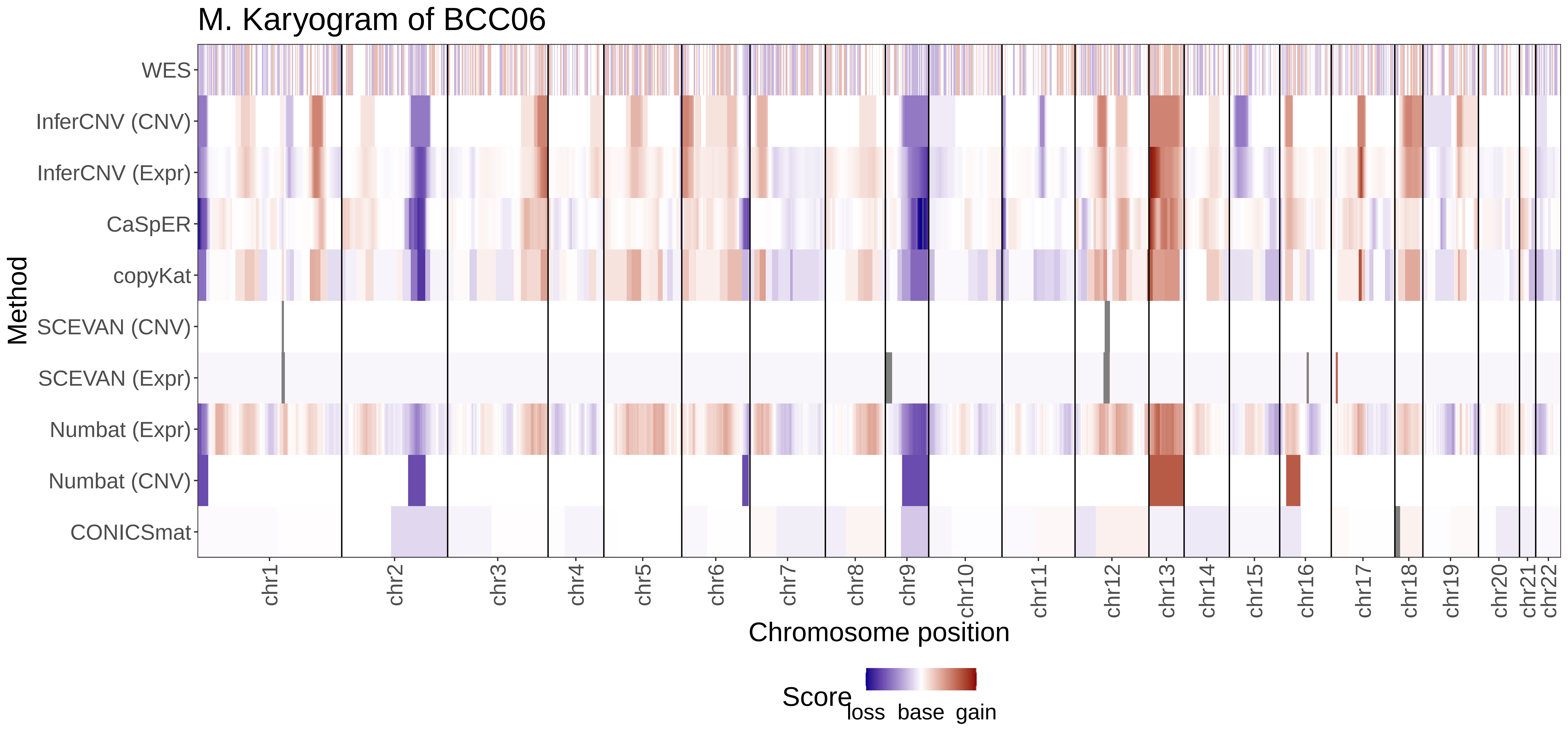

M. Karyogram of BCC06
Method
WES
InferCNV (CNV)
InferCNV (Expr)
CaSpER
copyKat
SCEVAN (CNV)
SCEVAN (Expr)
Numbat (Expr)
Numbat (CNV)
CONICSmat
chr1
chr2
chr3
chr4
chr5
chr6
chr7
chr8
chr9
chr10
chr11
chr12
chr13
chr14
chr15
chr16
chr17
chr18
chr19
chr20
chr21
chr22
Chromosome position
Score
loss
base
gain

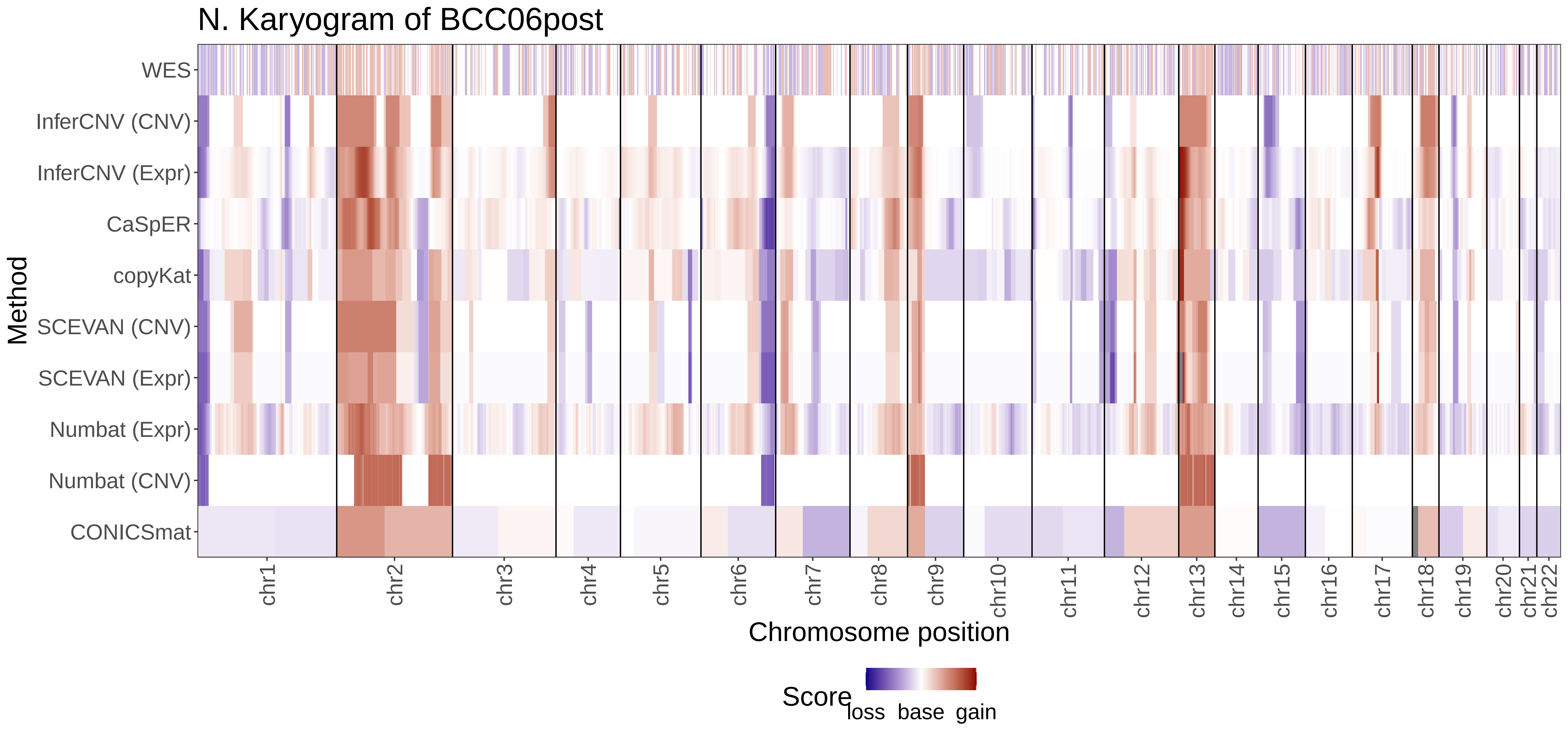
N. Karyogram of BCC06post
Method
WES
InferCNV (CNV)
InferCNV (Expr)
CaSpER
copyKat
SCEVAN (CNV)
SCEVAN (Expr)
Numbat (Expr)
Numbat (CNV)
CONICSmat
chr1
chr2
chr3
chr4
chr5
chr6
chr7
chr8
chr9
chr10
chr11
chr12
chr13
chr14
chr15
chr16
chr17
chr18
chr19
chr20
chr21
chr22
Chromosome position
Score
loss base gain
